# BBB-Permeable Near-Infrared Oxazine Fluorophores for White Matter Tract Imaging

**DOI:** 10.64898/2026.05.22.727303

**Authors:** Biruk Mengesha, Mingchong Dai, Antonio R. Montaño, Ge Huang, Stevie Huynh, Summer L. Gibbs, Lei G. Wang

## Abstract

Preserving critical functional tissue remains a major challenge in fluorescence-guided surgery (FGS), particularly in neurosurgery where injury to white matter tracts can lead to lasting neurological deficits. Probes for preventing iatrogenic injury of the peripheral nervous system have highlighted the potential of real-time intraoperative guidance for nerve-sparing surgery, supporting improved preservation of critical functional tissue, but analogous tools for the central nervous system remain limited. While there are some awake mapping and MRI based techniques that try to minimize the resection of white matter tracts (WMTs), they do not provide surgeons with real-time visualization. To address this gap, we rationally translated a peripheral nerve targeted oxazine fluorophore platform into a library of novel oxazine dyes that can penetrate the blood-brain barrier (BBB), exhibit specificity for white matter tissue and maintain spectral compatibility with clinically relevant imaging systems. Guided by our fluorophore medicinal chemistry platform, this work includes library design, photophysical property characterization, in vivo WMTs specificity screening, and in vivo imaging window assessment of the lead candidates to evaluate neurosurgical workflow compatibility. The lead candidate identified from this library demonstrated strong in vivo WMTs-specific fluorescence intensity, contrast and imaging window compatible with neurosurgical workflows, establishing BBB-permeable oxazine fluorophores as a promising scaffold for future structure-sparing neurosurgical FGS.

## 1. Introduction

White matter tracts (WMTs) are often at-risk during neurosurgery; this puts surgeons in a difficult position of balancing effective resection while preserving white matter bundles that support communication throughout the brain. Brain tumor resections, particularly glioblastoma surgery, are a prime example of this challenge with 347,992 new cases of brain cancer and 246,253 deaths globally reported as recently as 2019^1^. Evidence supports that a more extensive resection of the tumor can lead to more favorable outcomes for patient life expectancy^2,3^, however more aggressive resection can place eloquent regions of the brain, including critical WMTs at risk of injury^4^. Damage to these white matter bundles within the brain is associated with negative patient outcomes, for example, damage to the arcuate fasciculus is associated with deficits in speech fluency^5^, injury to the uncinate fasciculus and left inferior longitudinal fasciculus is associated with impairments in name retrieval^6,7^, and injury to the right superior longitudinal fasciculus is associated with deficits in spatial working memory^8^. The challenge of preserving these and many other critical structures within the brain has driven advances in preoperative and some intraoperative techniques, but these are not without shortcomings.

White matter tractography is a mapping technique used for reconstructing the WMTs prior to surgery, allowing surgeons to then plan the resection of tumor according to the patients’ anatomy^9^. The basis of this reconstruction is diffusion tensor imaging, a magnetic resonance imaging (MRI) technique which relies on the anisotropic diffusion of water molecules along coherently organized axonal fibers of white matter tracts^10^. A key limitation of preoperative techniques like this is intraoperative brain shift, which renders the preoperative mapping less reliable^11–13^. Another limitation is that intraoperative MRI-based imaging can add considerable time to the procedure^14^. Direct cortical stimulation during awake craniotomy is another critical intraoperative technique, where stimulation of the brain is used to map out regions that would be too detrimental to resect by asking patients questions during surgery^15^. This remains a powerful intraoperative technique^16^, but its utility is inherently limited by the need to keep the patient awake and able to participate throughout the procedure. Careful patient selection for awake craniotomies is therefore required, and the technique may fail when the patient must be converted to general anesthesia^17^. Collectively, these limitations highlight the need for complementary approaches that provide direct, real-time visualization and identification of WMTs during surgery.

Fluorescent-guided surgery (FGS) continues to expand at an unprecedented rate, with a growing number of probes under clinical evaluation, most commonly falling into the category of tumor highlighting agents^18^. FGS systems operate nearly exclusively in the near-infrared region (NIR, 650-900 nm), where tissue chromophore absorption, autofluorescence and scatter fall to local minima, permitting high-contrast and high-resolution imaging in vivo^19,20^. Notably, two NIR molecularly targeted probes (i.e., Cytalux and Lumisight) have recently received FDA approval for intraoperative cancer detection across major indications (e.g., ovarian, lung and breast)^21–24^. Despite this progress, most efforts in FGS have focused on diseased tissue identification and resection rather than preservation of the critical tissue structures (e.g., nerves, ureters, and WMTs)^18^. This broader push toward precision surgery extends to iatrogenic injury prevention, as reflected by the usage of NIR ICG as a vasculature perfusion imaging agent^25^. Probes of this nature have made significant progress as there are clinical trials for ureter- and nerve-specific probes underway ^18,26,27^. Building on our group’s previous success in developing NIR probes for sparing peripheral nerves^28–30^, and motivated by the clinical demand for improvements in WMTs imaging, we were well positioned to continue to push this space of probes for preventing iatrogenic injury in the CNS.

Motivated by the historic success of morpholine-containing compounds in central nervous system drugs^31^, and our group’s fluorophore medicinal chemistry experience with the NIR oxazine scaffold, we set out to develop a library of novel oxazine dyes designed for robust BBB permeability, strong WMTs specificity, and neurosurgical workflow compatibility (**Figure 1**). In this context, our goal was to translate a structure sparing fluorophore platform into BBB-permeable NIR small molecule fluorophores for future fluorescence-guided neurosurgery with improved functional outcomes. A small molecule probe with these properties built-in could provide a solution to address key limitations associated with current tools by providing real-time visualization during surgery, allowing brain shift to be readily accounted for intraoperatively and providing more versatility for patients in whom awake craniotomies are not feasible. Guided by this design strategy, the present study includes library design, photophysical characterization, in vivo WMTs specificity screening and imaging window assessment of lead candidates to assess compatibility with neurosurgical workflows.

**Figure 1:**
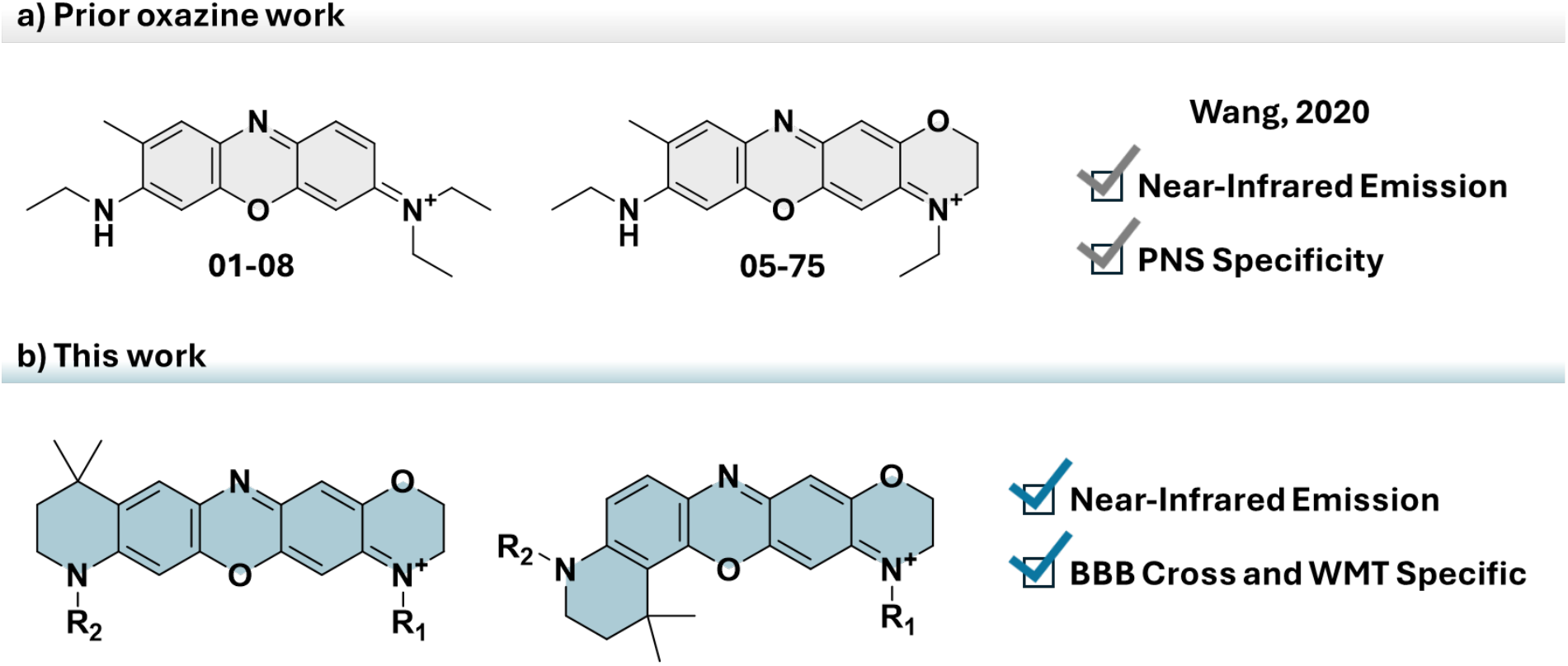
Translation of a peripheral nerve-specific oxazine fluorophore platform into BBB-permeable scaffold for white matter tract imaging. **(a)** Prior asymmetric oxazine fluorophores (LGW01-08, LGW05-75) developed for peripheral nerve imaging, established a blood-nerve barrier-permeable NIR fluorescent nerve-sparing fluorophore platform. **(b)** Rational scaffold engineering in the present work introduced tertiary/tertiary amine termini, asymmetric lipophilicity balance, and morpholino/tetrahydroquinoline features to support BBB permeability while preserving NIR fluorescence emission, yielding novel 700 nm CNS-targeted oxazine scaffolds for WMTs imaging.

## 2. Results

### 2.1. Design and Synthesis of a BBB-Permeable NIR Oxazine Fluorophore Library

Recent work on the oxazine scaffold for FGS, particularly for nerve-specific imaging, has provided important structure-activity relationship knowledge that guided the design of compounds in this library. Studies of oxazine 1 and oxazine 4 fluorophores^28,32^, and together with the matrix-designed library^30^ were pivotal in establishing the current understanding of peripheral nerve specificity. Lead candidates from these libraries are asymmetric and blood-nerve barrier (BNB) permeable. Although they were developed for a different target than this work, they motivated the use of asymmetric scaffolds in the present library for central nervous system (CNS) targeting. In addition to the trend regarding asymmetric fluorophores, fluorophores incorporating a fused benzomorpholino group in the matrix-designed library were noted as particularly bright fluorophores^30^.

Work outside of the field of the oxazine scaffold and instead on the xanthene scaffold also informed the design of these fluorophores, as the modified quinoline used in this library, a tetrahydroquinoline (THQ), in previous work has displayed excellent fluorescent properties^33^, which is believed to be due to its restriction in auxochrome rotation, similar to the pattern documented with fluorophores with a fused benzomorpholino group^30^. Accordingly, the dimethyl THQ was incorporated as a design element to help balance the physicochemical properties required for BBB permeability, as passive transport across the BBB is generally more restrictive for hydrophilic compounds^34^. Alongside the known utility of the morpholine-containing motifs in CNS-targeted drugs^31^, the more carbon-rich THQ-motif (i.e., dimethyl THQ) was therefore selected to tune scaffold lipophilicity while retaining a more hydrophilic morpholino containing auxochrome on the opposite side, creating an asymmetric amphiphilic scaffold architecture. Different substitution patterns on the dimethyl THQ group and fused benzomorpholino nitrogens were systematically examined to tune the physicochemical properties expected to play a key role in BBB permeability, as well as the WMT specificity and imaging window.

This library consists of 18 fluorophores, all including a fused benzomorpholino group and a dimethyl THQ group (**Figure 2a**). The cyclization reaction that produces the THQ moiety yields two products, which give rise to class A and B fluorophores (**Figure 2a**). Cyclization typically results in a 2:1 ratio, favoring the class A fluorophore precursor. Each compound within the classes then varies regarding the substitution patterns on the nitrogen of both the morpholino and THQ groups, the substitutions being a *N*-methyl, *N*-ethyl, or *N*-(2-methoxyethyl) group.

**Figure 2:**
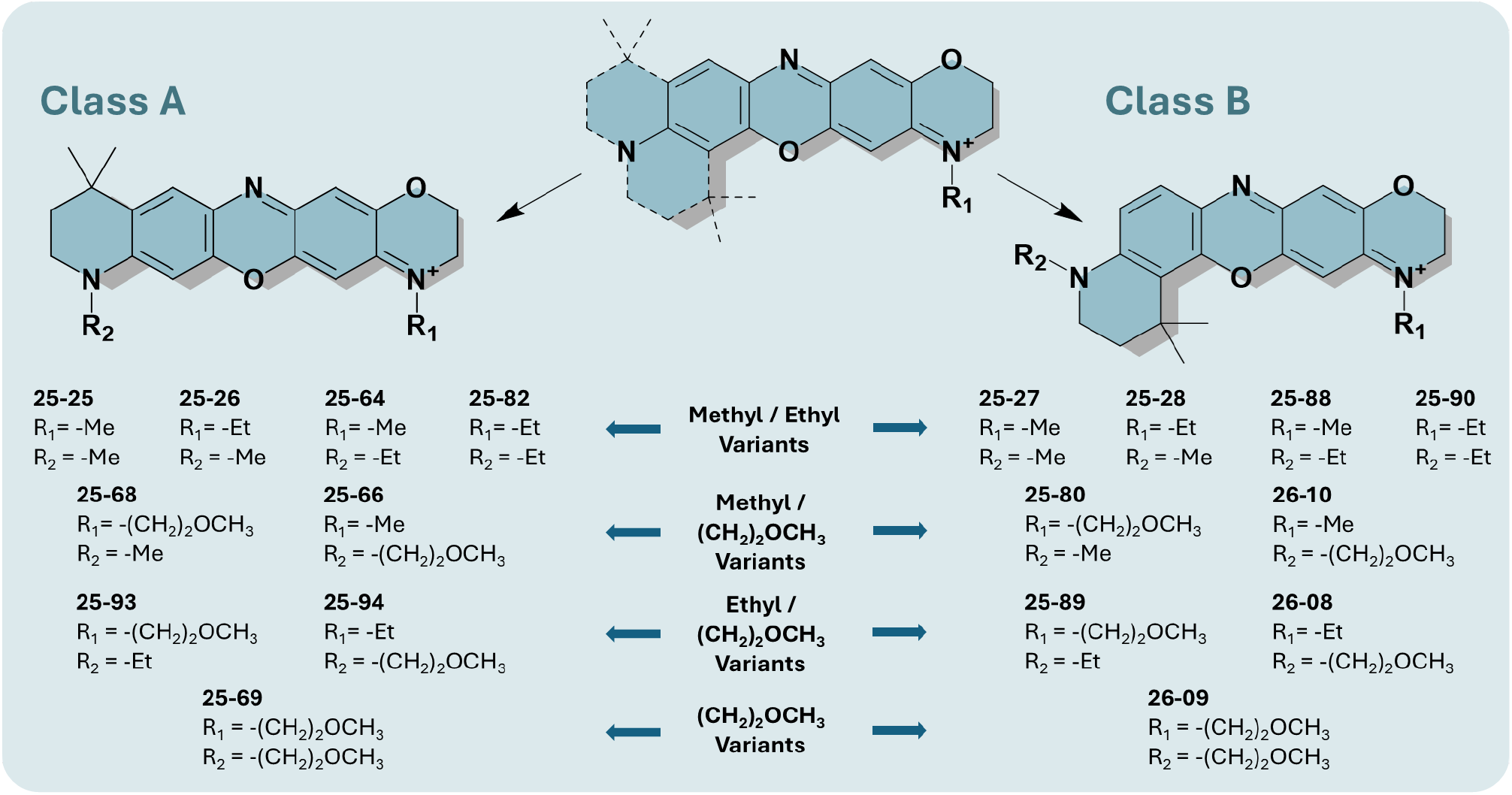
Design of the 700 nm BBB-permeable oxazine fluorophore library for WMTs imaging. General chemical structures for class A and class B fluorophores, representing all 18 fluorophores. The scaffold was engineered to introduce tertiary/tertiary amine termini and an asymmetric amphiphilic architecture through a morpholino containing side and a more hydrophobic dimethyl tetrahydroquinoline containing side, while auxochrome nitrogen substitution was varied to further tune BBB permeability and WMTs-specific imaging properties.

### 2.2. Photophysical Properties

The absorption and emission profiles of the compounds were collected in phosphate buffered saline (PBS) at a pH of 7.4, and these data were then used to determine the photophysical properties of the BBB-permeable fluorophore candidates (**Figure 3a**). All 18 fluorophores maximally absorb and emit within the NIR, with λ_abs,max_ values ranging from 652 to 666 nm and λ_em,max_ values ranging from 674 to 690 nm, placing the entire library within a clinically relevant imaging window for FGS^18^. Notably, class A fluorophores, the first 9 compounds consistently demonstrated substantially higher quantum yields and brightness than matched class B analogs, with quantum yields values ranging from 15.2 to 21.5% for class A versus 2.9% to 4.3% for class B, and brightness values ranging from 4200 M^-1^cm^-1^ to 10800 M^-1^cm^-1^ versus 700 M^-1^cm^-1^ to 1800 M^-1^cm^-1^ respectively. Molar extinction coefficient, however, did not display a consistent class-dependent trend. Class B fluorophores showed a greater bathochromic shift relative to class A analogs, with an average shift of 8 nm in absorption and 11 nm in fluorescence emission. Class matched compounds **25-25** and **25-27** provided representative analogs to compare spectral properties of each class with clinically relevant fluorophore (i.e., methylene blue). The normalized absorption and emission profiles for these two fluorophores are spectrally similar to methylene blue (**Figure 3b-c**), with **25-27** showing closest spectral overlap due to its slight red shift, making these fluorophore candidates well aligned with clinically relevant NIR 700 nm imaging channels.

**Figure 3:**
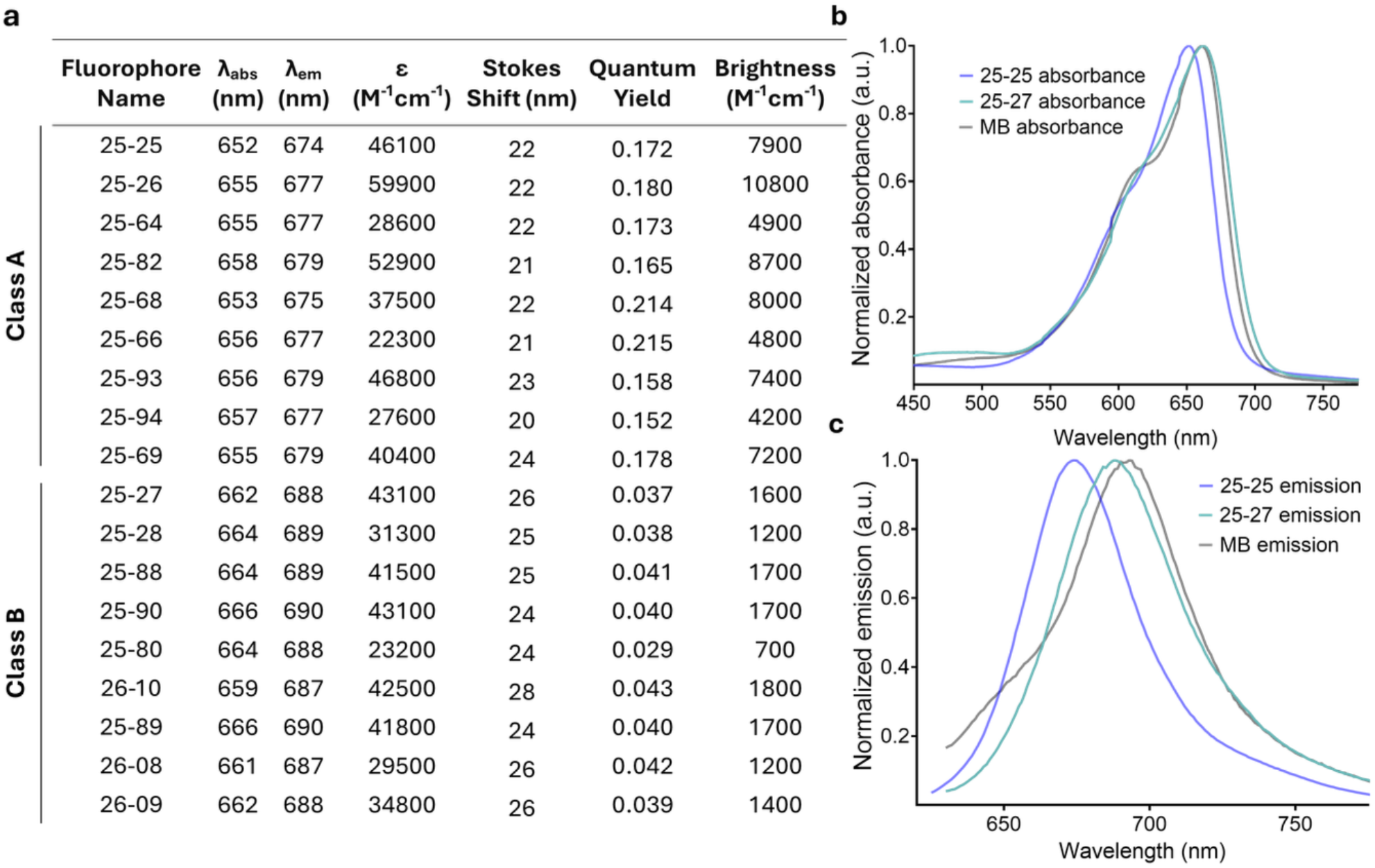
Photophysical characterization of the oxazine fluorophore library. **(a)** Tabulated photophysical properties of library, including wavelength of absorption and emission maximum, molar extinction coefficient, quantum yield and brightness, measured in phosphate buffered saline. **(b)** Normalized absorption spectra of **25-25, 25-27**, and the 700 nm clinically relevant analog, methylene blue. **(c)** Normalized emission spectra of **25-25, 25-27**, and the 700 nm clinically relevant analog, methylene blue.

### 2.3. In Vivo White Matter Specificity Screening

The initial in vivo WMTs specificity and intensity screening of this library followed a protocol where mouse brains were excised 4 hours after intravenous administration of a fluorophore candidate, sectioned to expose the corpus callosum and imaged (**Figure 4a**). To avoid selection bias, white light images were used to select regions of interest (ROIs), composed of white matter and gray matter tissue, both sides of the corpus callosum representing the white matter and the region of cortex above the WMT ROI representing the gray matter (**Figure 4a**), those ROIs were then used to quantify fluorescence intensity from the corresponding fluorescence image. Quantified intensities were subsequently used to calculate the white-matter-to-gray-matter signal-to-background ratio (SBR), which was used as a measure of WMTs-specific contrast.

**Figure 4:**
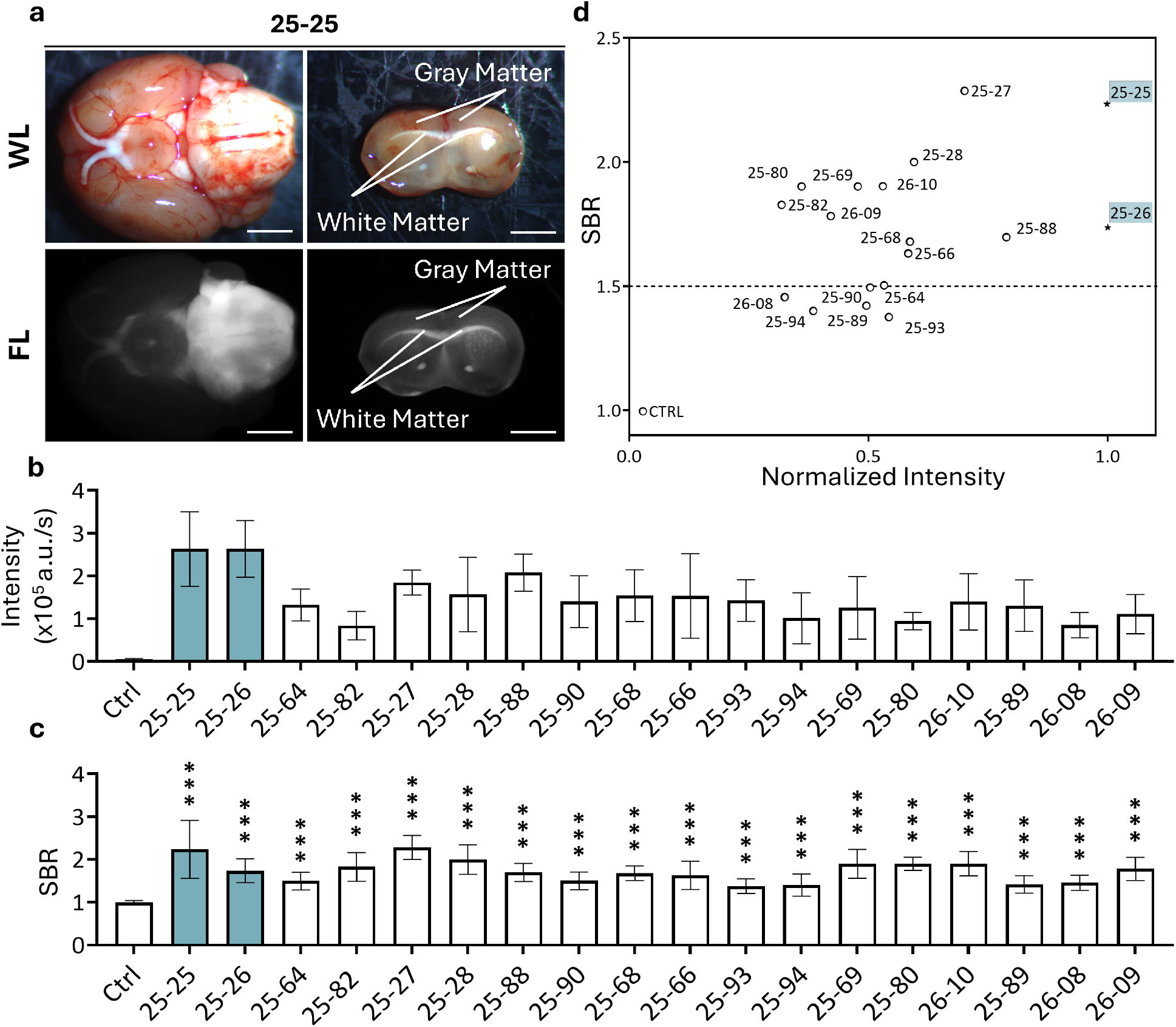
In vivo white matter tract specificity and intensity screening of the BBB-permeable oxazine fluorophore library. **(a)** Representative white light and fluorescence images of the whole brain and coronal after administration of **25-25**, images are displayed with autocontrast adjustment for visualization. Labels present on coronal section images are for gray and white matter with 2.5 mm scale bars. **(b)** Average white matter tract in vivo intensity for all 18 fluorophores and a 4-hour timepoint vehicle-injected control following intravenous administration, n=4. **(c)** Average white-matter-to-gray-matter tissue intensity ratios (SBRs) for all 18 fluorophores and a 4-hour timepoint vehicle-injected control following intravenous administration, n=4. Average SBRs were compared to the vehicle-injected control with a one-way ANOVA using Geisser-Greenhouse correction followed by a Fisher’s LSD multiple comparison test, where 1, 2, 3 and 4 asterisks represent P values less than 0.05, 0.01, 0.001, and 0.0001 respectively. **(d)** Scatter plot of average white-matter-to-gray-matter tissue intensity ratio against the average normalized in vivo intensity for all 18 fluorophores and a vehicle injected control following intravenous administration.

Intensity values (a.u./s) varied greatly within this library, with a 10-fold difference between the brightest and dimmest in vivo fluorescence intensities seen. The three brightest fluorophores in this library were seen in **25-26, 25-25** and **25-88** with intensity values of 2.64 × 10^5^, 2.63 × 10^5^, and 2.07 × 10^5^ a.u./s respectively (**Figure 4b**). When it comes to specificity, **25-27, 25-25** and **25-28** all had the highest contrast, with each fluorophore candidate having an SBR equal to or greater than 2.00, at 2.28, 2.23, and 2.00 respectively (**Figure 4c)**.

When considering both metrics, **25-25** and **25-26** emerged as lead candidates, as they are the brightest fluorophores in tissues in this library. These two fluorophores, **25-25** and **25-26**, not only displayed exceptional brightness but also met and surpassed a threshold for SBR of 1.5, which has been previously noted as the threshold acceptable for FGS contrast^35^. Based on their combination of strong brightness and contrast, **25-25** and **25-26** were advanced for in vivo imaging window evaluation to assess compatibility with neurosurgical workflows.

### 2.4 Lead Candidate In Vivo Imaging Window Profiling

To determine the optimal imaging window of this library’s lead candidates, the method used for screening fluorophores was modified to include additional timepoints. This resulted in the time dependent assessment of each lead fluorophore’s intensity and contrast over a span of 24 hours, including 0.5-, 1-, 2-, 4-, 8-, 16-, and 24-hour timepoints (**Figure 5**).

**Figure 5:**
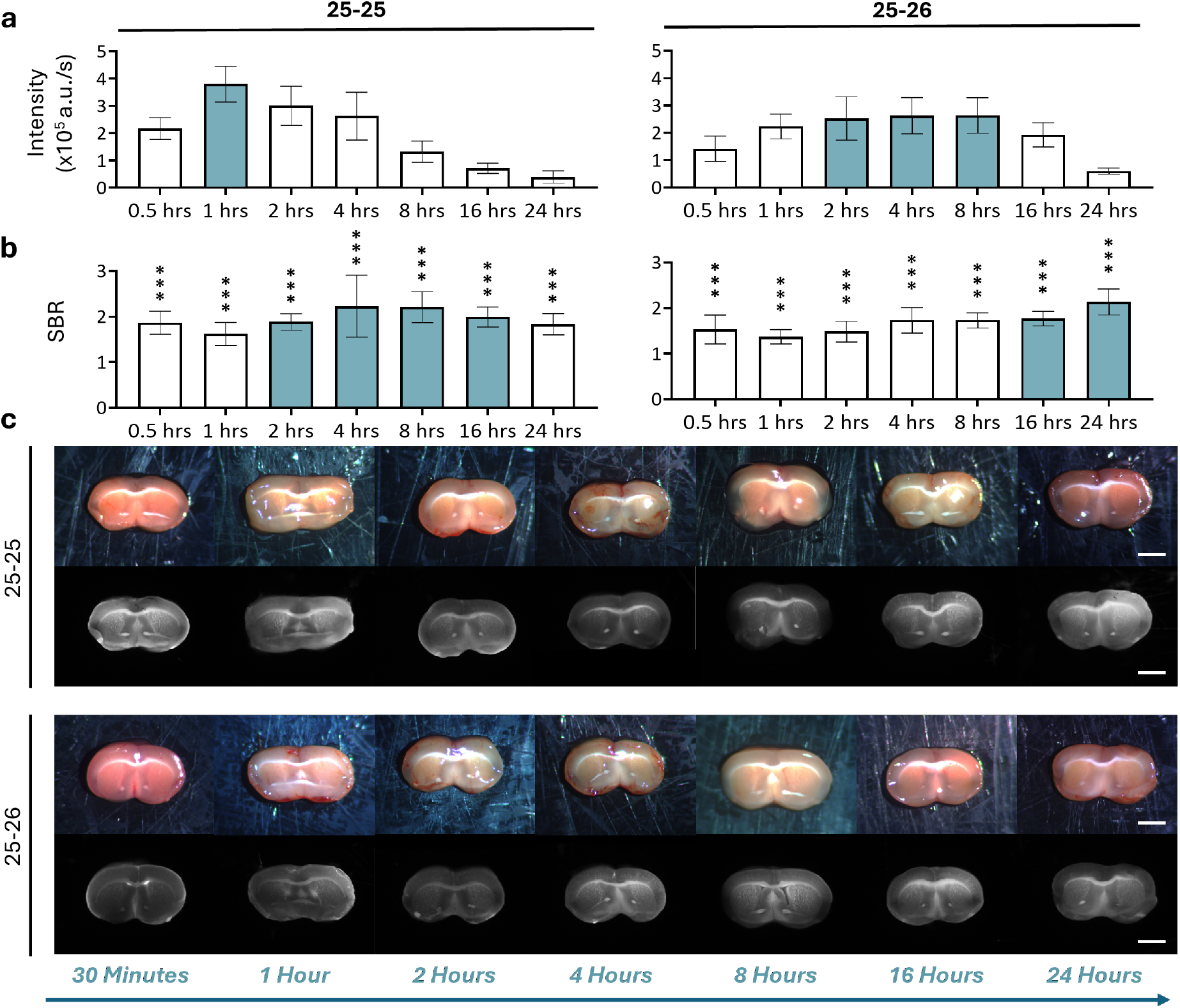
In vivo imaging window assessment of lead fluorophores 25-25 and 25-26. **(a)** Average WMTs in vivo intensity for lead fluorophores after various timepoints following intravenous administration, n=4. **(b)** Average white-matter-to-gray-matter tissue intensity ratios for lead fluorophores at different timepoints following intravenous administration, n=4. Average SBRs were compared to a 4-hour timepoint vehicle-injected control with a one-way ANOVA using Geisser-Greenhouse correction followed by a Fisher’s LSD multiple comparison test, where 1, 2, 3 and 4 asterisks represent P values less than 0.05, 0.01, 0.001, and 0.0001 respectively. **(c)** Representative white light and fluorescence images of the coronal section at various timepoints for lead fluorophores, images are displayed with autocontrast adjustment for visualization.

Each lead candidate displayed a unique intensity profile. **25-25** reached a maximum intensity within 1 hour of injection at 3.79 × 10^5^ a.u./s, subsequent timepoints showed progressively decreasing fluorescence intensity ending at 0.39 × 10^5^ a.u./s (**Figure 5a**). On the other hand, **25-26** fluorescence intensity increased up to 8 hours reaching 2.64 × 10^5^ a.u./s. After 8 hours fluorescence intensity decreased steadily, reaching 0.61 × 10^5^ a.u./s at the final time point (**Figure 5a**).

Similarly, the contrast profile also varied between lead fluorophores. **25-25** throughout the in vivo imaging window testing exceeded the threshold SBR of 1.50, reached maximum contrast at 4 hours with a SBR of 2.23 and then continually dropped until the end of the study with a SBR of 1.83 (**Figure 5b**). **25-26** also exceeded the SBR threshold of 1.50 for all timepoints except 1 and 2 hours where it fell to 1.37 and 1.49 respectively but then rebounded to above the threshold at 4 hours to 1.73. It then continually increased during the study, reaching its maximum SBR of 2.14 at 24 hours (**Figure 5b**).

## 3. Discussion

The NIR oxazine core has emerged as an outstanding small-molecule probe scaffold for peripheral nerve-sparing FGS^28–30^. Here, this fluorophore scaffold is extended to a new imaging application, in vivo WMTs imaging in the CNS. This effort was enabled by our fluorophore medicinal chemistry platform, which uses prior structure-activity relationship knowledge to systematically tune features relevant to surgical imaging, including tissue specificity, BNB/BBB permeability, brightness, and photophysical properties. As a result, the fluorophore candidates described here combine small molecule design, structure tunability, NIR emission profile, and high brightness, all of which support continued translational development.

The design was based on the prior success of the benzomorpholino group on oxazine dyes, which has been used to produce bright fluorophores while retaining specificity for peripheral nerve tissue^30^ and together with the success of morpholino groups in CNS drugs^31^. In the present scaffold, the morpholino-containing auxochrome was also expected to contribute a more hydrophilic character whereas the incorporation of the THQ motif and tertiary amine substitution on the opposite side increased lipophilic character. This asymmetric distribution of polarity may create an amphiphilic fluorophore scaffold that is better suited for BBB penetration while still supporting selective accumulation in lipid rich WMTs. In addition, variation of the auxochrome nitrogen substituents, specifically *N*-methyl, *N*-ethyl, or *N*-(2-methoxyethyl) group, provided a tractable way to further tune the physicochemical balance of the oxazine scaffold. We hypothesized that this tuning contributed not only to BBB permeability, but also to differences in WMTs specificity as well as the time dependent in vivo imaging windows observed across the library.

The two classes within the scaffold, class A and B (**Figure 2a**), differed in two main ways, the quantum yield and the λ_abs/ems_. The class A compounds had greater quantum yields than the class B dyes (**Figure 3a**). Class B fluorophores displayed a slight red shift compared to class A (**Figure 3a-c**), with an average bathochromic shift of 11 nm. The emission profile is also critical for FGS, as fluorophores that emit in the NIR are able to avoid both the significant autofluorescence and scattering associated with visible light^19,20^. Notably, fluorophores in this library produced spectral profiles similar to methylene blue, a clinically used and FDA-approved dye, and were well aligned with clinically relevant 700 nm imaging channels for FGS.

In vivo WMTs-specific fluorescence intensity and contrast also varied substantially across the oxazine library. Although several compounds demonstrated strong WMTs contrast, and others showed high fluorescence intensity in the WMTs, selection of the lead candidates required balancing NIR spectral profile, in vivo WMTs-specific brightness and contrast. In this context, **25-25** and **25-26** combined strong in vivo fluorescence intensity with favorable WMT contrast, making them the most compelling lead candidates for further evaluation.

Subsequent in vivo imaging window assessment revealed unique profiles for these two lead candidates. **25-25** reached peak in vivo fluorescence intensity earlier and provided strong WMT-specific signal within a shorter time frame, whereas **25-26** showed a more sustained contrast profile over extended time points. Primary brain tumor resections typically last 2 to 4 hours^16,36^, but depending on case complexity, procedures can extend to 8 hours ^37^. Within this context, **25-25** may be better suited for shorter procedures that benefit from strong early WMT-specific signal, whereas **25-26** may be advantageous in surgeries that require more prolonged WMT contrast. Together, these findings suggest that successful BBB-permeable fluorophores for CNS structure imaging may not simply require increased lipophilicity, but also a balanced amphiphilic architecture that supports BBB penetration, WMTs-targeted accumulation and imaging windows compatible with neurosurgical workflows.

Further preclinical in vivo studies will be needed to better define their translational potential, which include dose-ranging studies, biodistribution analysis, pharmacokinetic profiling and toxicology testing. These studies will help define the optimal imaging dose, safety margin and translational feasibility of the lead candidates. Future directions also include application for two-color imaging that highlights both white matter and tumor tissue, something that has been successfully done with prior generations of oxazine probes in the peripheral nervous system^38^. In this context, **25-26** proves particularly interesting, its delayed contrast may be well suited for pairing with tumor-targeted imaging agents that also benefit from later imaging time points. ICG has demonstrated excellent contrast when administered in mice models and in human patients due to the enhanced permeability and retention (EPR) effect, with optimal contrast found at 24 hours^39^. This timepoint is also the time when **25-26** reaches its maximum contrast, permitting potential for coadministration of **25-26** and ICG 24-hour prior to obtain two-color images that demonstrate contrast for white matter versus gray matter and tumor versus healthy tissue.

The field of FGS continues to expand at an unprecedented pace, but much of the current translational pipeline remains focused on diseased tissue identification rather than preservation of critical structures such as nerves and white matter tracts. This work extends the FGS toolbox from the peripheral nervous system into the central nervous system and demonstrates the oxazine fluorophore scaffold as a promising platform for WMT imaging. The lead candidates identified in this library demonstrated favorable NIR fluorescence, strong WMT-specific intensity and contrast, and in vivo imaging windows compatible with neurosurgical workflows, supporting continued development of BBB-permeable oxazine fluorophores toward future clinical translation.

## 4. Materials and Methods

### 4.1 Study overview

This project was conducted with the goal of developing NIR fluorescent probes with inherent white matter specificity and suitable characteristics for imaging systems already in clinics. This project’s library spans 18 fluorophores all based on an oxazine scaffold and incorporates a morpholino and dimethyl tetrahydroquinoline (THQ) into this scaffold. Compounds’ structural identity and purity were assessed using high-performance liquid chromatography-mass spectrometry (HPLC-MS). Optical properties including absorption and emission spectra, molar extinction coefficient, and quantum yield were quantified using a fluorescence spectrometer. Fluorophores were administered intravenously and imaged on a custom-built imaging system. Regions of interest (ROIs), white and gray matter, were then quantified with a custom MATLAB code, and quantified intensity values were then used to find the signal to background ratio. Synthetic details, optical data, and other data related to characterizations of the compounds can be found in the Supplementary Materials.

### 4.2 Oxazine library synthesis and characterization

All chemical building blocks were purchased from Sigma-Aldrich (Burlington, MA, USA), TCI (Tokyo, Japan) or Ambeed (Buffalo Groove, IL, USA). Unless otherwise indicated, all commercially available starting materials were used directly without further purification. If starting materials were not available, they were synthesized according to the protocols described here (see Supplementary Materials) or based on literature methods. All 18 compounds were synthesized with two intermediates via a condensation reaction, one being a tetrahydroquinoline derivative and the other being a morpholino derivative. The cyclization of the former results in two products which form the two classes of fluorophores presented here, each fluorophore within the classes then varies in *N*-methyl, *N*-ethyl, *N*-(2-methoxyethyl) substitution on the THQ and fused benzomorpholino. Analytical thin-layer chromatography was done using silica gel 60 (10-12 µm, MilliporeSigma, Burlington, MA, USA). Crude reaction product was adsorbed on 60Å, 40-63 µm silica gel (Sorbent Technologies, Inc., Norcross, GA, USA) and purified using a Biotage Selekt automated flash chromatography system on Biotage Sfär column (Biotage, Uppsala, Sweden). Mass spectra were collected using an Agilent 1260 Infinity HPLC System and LC/MSD single quadrupole system (Agilent Technologies, Santa Clara, CA, USA). Each compound also had its absorption spectra collected in phosphate buffered saline (PBS) and emission profiles collected (also in PBS) with an excitation of either 600 nm or 610 nm on a SpectraMax M5 spectrometer (Molecular Devices, San Jose, CA, USA). Molar extinction coefficient and relative quantum yield were calculated from these spectral data.

### 4.3 Animals

The use of animals in this study was approved by the Institutional Animal Care and Use Committees (IACUC) at Oregon Health and Science University (OHSU). All animals used in this study were male and female CD-1 mice, 3-5 weeks in age, purchased from Charles River Laboratories (Wilmington, MA, USA), mice were euthanized prior to imaging using inhalation of carbon dioxide followed by cervical dislocation. All mice were fed chlorophyll-free diet (Animal Specialties, Quakertown, PA, USA) for at least seven days prior to *in vivo* imaging studies. Inhalation of carbon dioxide was used as the primary method of euthanasia followed by cervical dislocation as a secondary method in all mice. Euthanasia was confirmed by physical examination to ensure cessation of heartbeat and respiration and is consistent with the recommendations of the Panel on Euthanasia of the American Veterinary Medical Association.

### 4.4 In vivo white matter screening

Fluorophores were screened to determine the in vivo intensity and specificity in CD-1 mice with a 50/50 male/female sex distribution. Fluorophores were solubilized based on a previously published formulation^40^, where fluorophores were dissolved in 5% Kolliphor® EL, 10% dimethyl sulfoxide (DMSO), 65% serum, and 20% phosphate buffered saline (PBS) at a concentration of 2.5 mM. All fluorophores were administered intravenously (I.V.), using the tail vein as the site of injection and dosed at 7.5 µmol/kg. All fluorophores were screened at 4 hours to find potential lead candidates, and lead fluorophores were screened at additional timepoints (30 minutes, 1, 2, 8, 16 and 24 hours). Vehicle injected mice were used as controls. All reported in vivo intensity and SBR data were screened at n=4 mice. After mice were euthanized using carbon dioxide and cervical dislocation, intact brains were excised and imaged. After imaging, coronal sections were made that divided the brain into equal thirds. The posterior face of the first section and the anterior face of the second section were imaged.

### 4.5 Intraoperative fluorescence imaging systems

A macroscopic imaging system was custom built for small-animal imaging and was used for real-time color and fluorescence imaging of mice in this project. The imaging system consists of Lumencor Spectra Light Engine (Lumencor, Beaverton, OR, USA) as the light source, directed by a liquid light guide. White light images used 438, 555, 660 nm light and fluorescence images used 630 nm light. Fluorescence images were collected with an ET700/75m filter (Chroma Technology Corporation, Rockingham, VT, USA), which is a 700 ± 37.5 nm bandpass emission filter, and white light images were collected without an emission filter. Images were captured with a QImaging EXi Blue monochrome camera (Teledyne QImaging, Surrey, Canada) outfitted with a removable Bayer filter for co-registered color image collection. Images were taken with exposure times ranging from 50 ms to 750 ms. All images were taken under the same conditions, allowing for a quantitative comparison of in vivo intensities.

### 4.6 Analysis of intraoperative white matter contrast and intensity

Using a custom MATLAB code (doi: 10.5281/zenodo.3698316), tissue intensities could be quantified, and specificity could be assessed. White light and fluorescence images were co-registered to select regions of interest (ROIs) based on the white light images to keep the user blinded to the fluorescent signal; ROIs included white matter tissue, using the left and right side of the corpus callosum (CC), and gray matter tissue, using the region of the cortex directly above the left and right sides of the CC. A background ROI was also collected above the entirety of the brain. The intensity values were reported for the white matter tissue and were divided by the exposure time to normalize data, resulting in measurements in intensity units per second. Reported intensities were averages of both left and right CC ROIs with a background correction for all mice (n=4). Signal to background (SBR) ratio was calculated from the ratio of background corrected white matter tissue to background corrected gray matter intensities. Reported SBRs were averages of all mice (n=4).

### 4.7 Statistical analysis

Statistical analysis was completed for SBRs using one-way analysis of variance (RM one-way ANOVA) with no assumption of sphericity (using the Geisser-Greenhouse correction) followed by a Fisher’s least significant difference (LSD) between the mean of the 4 hour timepoint vehicle-injected mice’s SBR against the SBR values collected for the initial screening and in vivo imaging window results with an α-value of 0.05. All column charts are presented as the mean with ± SD. All plots and statistical analyses were done using Prism (GraphPad, Boston, MA, USA).

## Acknowledgements

We would like to thank William Greer for his assistance with data analysis. This work was funded by the National Cancer Institute (R01CA271532, Gibbs) and the OHSU Foundation (Gibbs).

