## Supplementary Materials for "BBB-Permeable Near-Infrared Oxazine Fluorophores for White Matter Tract Imaging"

Oregon Health & Science University
Collaborative Life Sciences Building
2730 S Moody Ave
Portland, OR 97201

**This PDF file includes:**

Materials and Methods

Figure S1-14. HPLC-MS of intermediate compounds

Figure S15. Oxazine derivative library chemical structures

Figure S16-33. HPLC-MS and purity analysis of oxazine dyes

Figure S34-51. NIR oxazine dye spectra

### Materials and Methods

#### Oxazine library synthetic overview:

Key synthetic steps included alkylation of a m-anisidine with 1-bromo-3-methylbut-2-ene followed by cyclization with methanesulfonic acid. With each regioisomer then being used to form a diazonium intermediate in hydrogen chloride and 4-nitrobenzenediazonium tetrafluoroborate or being demethylated using boron tribromide. Each dye was formed using an aromatic amine and diazonium intermediate with acetic acid as the solvent and catalyst under 110 °C. Steps that involved purification used 60A, 40-63um silica gel (Sorbent Technologies, Inc., Norcross, GA, USA) to adsorb compound onto, and then was purified using a Biotage Selekt Flash Chromatography System (Biotage, Uppsala, Sweden) and fractions were assessed using analytical thin-layer chromatography using silica gel 60 (10-12 µM, MilliporeSigma, Burlington, MA).

#### Oxazine library structure and purity overview:

Each compound's mass-to-charge (m/z) ratio was used to assess purity, this was collected using an Agilent 1260 Infinity High Performance Liquid Chromatography System and LC/MSD single quadrupole system (Agilent Technologies, Santa Clara, CA). Each oxazine derivative was injected (10 µL in a 1:1 ratio of acetonitrile (0.1% formic acid) and water (0.1% formic acid) onto a C18 column (Poroshell 120, 4.6 x 50 mm, 2.7 micron, Agilent Technolgoies) and eluted with a solvent system of solvent A (water, 0.1% formic acid) and solvent B (acetonitrile, 0.1% formic acid).

#### Full synthetic details of the 18 oxazine fluorophores

#### General Reactions:

#### General Protocol A: 1-bromo-3-methylbut-2-ene alkylation.

An m-anisidine derivative (1 mmol) and potassium carbonate (2 mmol) was dissolved in anhydrous acetonitrile (5 mL) under $N_2$ and stirred at room temperature for 10 minutes. Then 1-bromo-3-methylbut-2-ene (1.3 mmol) was added under $N_2$ and was stirred at 50 °C overnight. When the m-anisidine derivative was consumed monitored on thin layer chromatography (TLC); the solution was evaporated and ran through a vacuum filtration using a celite filter. The filtrate was then evaporated with silica gel. With the compound adsorbed to the silica it was then purified using a flash chromatography system with a mobile phase of ethyl acetate/hexanes, on a 0-15% gradient to eluent the product.

#### General Protocol B: Methanesulfonic acid facilitated cyclization.

A m-anisidine derivative (1 mmol) was dissolved methanesulfonic acid (1 mL). Once added onto a water jacket condenser the solution was heated to 90 °C and allowed to stir for 30 minutes. The solution was quenched with periodic additions of potassium carbonate at 0 °C until the pH was 7. An extraction was then performed with dichloromethane and water. Silica was added into the organic phase and dried to be purified with a flash chromatography system using a mobile phase of ethyl acetate/hexanes, on a 0-10% gradient.

**General Protocol C: Diazonium intermediate formation.**

An aniline derivative (1 mmol) was dissolved in 1 part methanol (2 mL) then at 0 °C, 5 parts 2 molar hydrochloric acid solution (10 mL). While at 0 °C 4-nitrobenzenediazonium tetrafluoroborate (1.05 mmol) in four portions over the course of 10 minutes. The solution was stirred for 1 hour at 0 °C. Leaving a light to dark red slurry that was collected into a funnel and air dried overnight without further purification. If filtration is not possible, the solution was then quenched with a saturated sodium bicarbonate solution and extracted with dichloromethane and water and was used without further purification.

**General Protocol D: Anisole demethylation.**

A methoxylated m-anisidine (1 mmol) was dissolved in anhydrous dichloromethane (10 mL) under $N_2$ and stirred in a -78 °C acetone and dry ice bath for 30 minutes. A 1M boron tribromide (3 mmol) was added dropwise at -78 °C. Once completed the solution was allowed to stir for overnight. . Reaction progress was monitored with LC-MS. Once the aniline was consumed, the solution was quenched with water at 0 °C, and pH was adjusted to 6-7 using $K_2CO_3$. The solution was then extracted with dichloromethane and water. Silica was added into the organic phase and dried to be purified with a flash chromatography system using a mobile phase of ethyl acetate/hexanes on a 0-15% gradient.

**General Protocol E: Oxazine fluorophore condensation using Acetic Acid.**

An aniline derivative (0.13 or 0.1 mmol) and a diazonium intermediate (0.1 mmol) were dissolved in acetic acid (2 mL). The solution was then heated to 110 °C for 10 to 30 minutes, going from a vibrant red to a dark green to blue solution. The solution was then evaporated silica gel. The compound adsorbed with the silica was then purified with a flash chromatography system using a mobile phase of methanol/acetone, both with a 0.5% formic acid additive, on a 0-25% gradient to eluent the product.

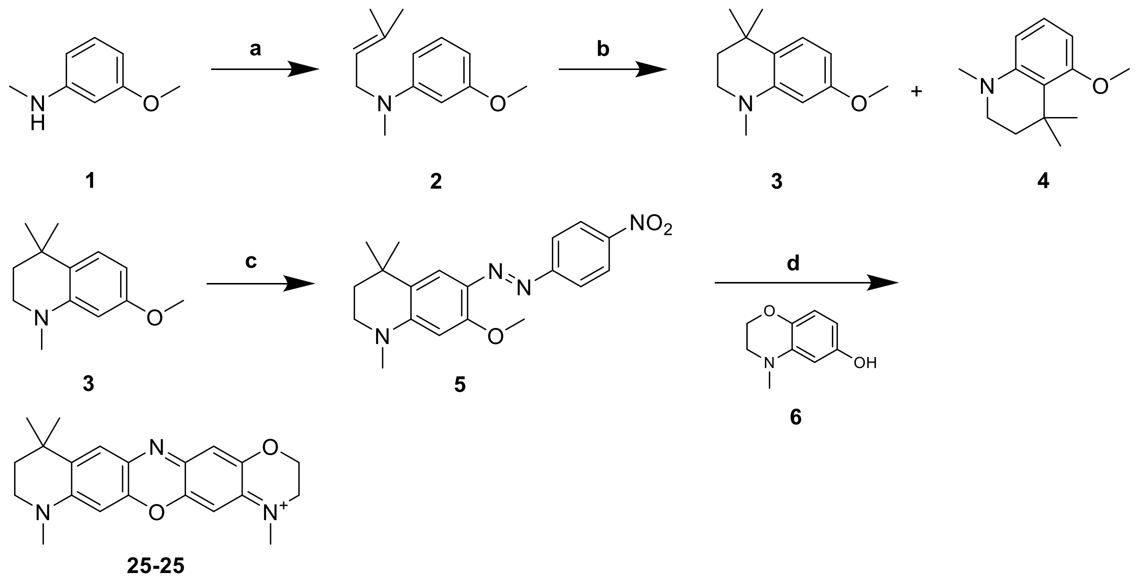


**Scheme 1:** Synthetic route to **25-25**. Reagents and conditions: a) 1-bromo-3-methylbut-2-ene, $K_2CO_3$, acetonitrile, 80 °C; b) methanesulfonic acid, 80 °C; c ) (i) 2 M hydrochloric acid / methanol (5:1), p-nitrobenzenediazonium tetrafluoroborate, 0 °C; (ii) $K_2CO_3$, 0 °C; d) acetic acid, 110 °C.

Compound **6** were synthesized following protocols reported by Montaño[1].

**3-methoxy-N-methyl-N-(3-methylbut-2-en-1-yl)aniline (2):** Compound **2** was prepared following **General protocol A** using compound **1** (100.0 mg, 0.728 mmol). Compound **2** was obtained (75.7 mg, 51%) as a pale-yellow oil. m/z: $(M+H)^+$ calculated for $C_{13}H_{20}NO^+$ 206.15, found 206.1 (ESI).

**7-methoxy-1,4,4-trimethyl-1,2,3,4-tetrahydroquinoline (3):** Compound **3** was prepared following **General protocol B** using compound **2** (52.0 mg, 0.253 mmol). Compound **3** was obtained (16.4mg, 32%) as an amber oil. m/z: $(M+H)^+$ calculated for $C_{13}H_{20}NO^+$ 206.15, found 206.0 (ESI).

**5-methoxy-1,4,4-trimethyl-1,2,3,4-tetrahydroquinoline (4):** Compound **4** was prepared following **General protocol B** using compound **2** (52.0 mg, 0.253 mmol). Compound **4** was obtained (8.7 mg, 17%) as an amber oil. m/z: $(M+H)^+$ calculated for $C_{13}H_{20}NO^+$ 206.15, found 206.0 (ESI).

**(E)-7-methoxy-1,4,4-trimethyl-6-((4-nitrophenyl)diazenyl)-1,2,3,4-tetrahydroquinoline (5):** Compound **5** was prepared following **General protocol C** using compound **3** (250 mg, 1.22 mmol). Compound **4** was obtained as red solid, which was used for the next step without further purification.

**4,8,11,11-tetramethyl-2,3,8,9,10,11-hexahydro-[1,4]oxazino[2,3-*b*]pyrido[2,3-*i*]phenoxazin-4-ium (25-25):** Compound **25-25** was prepared following **General protocol E** using compounds **5** (53.6 mg, 0.151 mmol) and **6** (25 mg, 0.151 mmol). Compound **25-25** was obtained (15.9 mg, 27%) as a blue solid. m/z: $(M+H)^+$ calculated for $C_{21}H_{24}N_3O_2^+$ 350.19, found 350.1 (ESI).

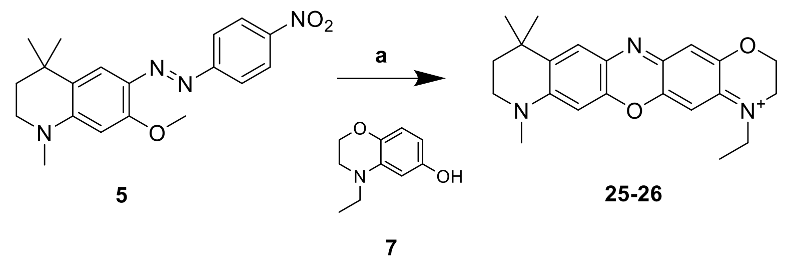


**Scheme 2:** Synthetic route to **25-26**. Reagents and conditions: a) acetic acid, 110 °C.

Compound **7** were synthesized following protocols reported by Wang[2].

**4-ethyl-8,11,11-trimethyl-2,3,8,9,10,11-hexahydro-[1,4]oxazino[2,3-*b*]pyrido[2,3-*i*]phenoxazin-4-ium (25-26):** Compound **25-26** was prepared following **General protocol E** using compounds **5** (49.4 mg, 0.139 mmol) and **7** (25.0 mg, 0.139 mmol). Compound **25-26** was obtained (19.1mg, 45%) as a blue solid. m/z: $(M+H)^+$ calculated for $C_{22}H_{26}N_3O_2^+$ 364.20, found 364.2 (ESI).

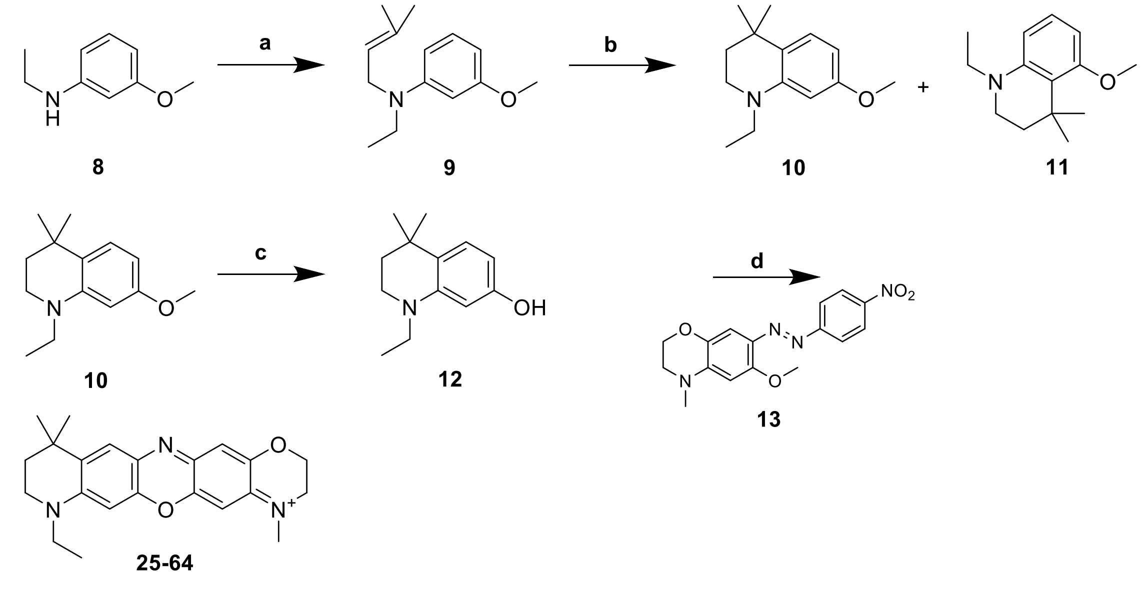


**Scheme 3:** Synthetic route to **25-64**. Reagents and conditions: a) 1-bromo-3-methylbut-2-ene, $K_2CO_3$, acetonitrile, 80 °C; b) methanesulfonic acid, 80 °C; c ) 1 M boron tribromide, dichloromethane, 0 °C to rt; d) acetic acid, 110 °C.

Compound **13** were synthesized following protocols reported by Montaño[1].

**N-ethyl-3-methoxy-N-(3-methylbut-2-en-1-yl)aniline (9):** Compound **9** was prepared following **General protocol A** using compound 8 (551.1 mg, 3.38 mmol). Compound **9** was obtained (741.3 mg, 60%) as a pale-yellow oil. m/z: $(M+H)^+$ calculated for $C_{14}H_{22}NO^+$ 220.17, found 220.1 (ESI).

**1-ethyl-7-methoxy-4,4-dimethyl-1,2,3,4-tetrahydroquinoline (10):** Compound **10** was prepared following **General protocol B** using compound **9** (389.0 mg, 1.78 mmol). Compound **10** was obtained (185.4 mg, 48%) as an amber oil. m/z: $(M+H)^+$ calculated for $C_{14}H_{22}NO^+$ 220.17, found 220.1 (ESI).

**1-ethyl-5-methoxy-4,4-dimethyl-1,2,3,4-tetrahydroquinoline (11):** Compound **11** was prepared following **General protocol B** using compound **9** (389.0 mg, 1.78 mmol). Compound **11** was obtained (73.1 mg, 19%) as an amber oil. m/z: $(M+H)^+$ calculated for $C_{14}H_{22}NO^+$ 220.17, found 220.1 (ESI).

**1-ethyl-4,4-dimethyl-1,2,3,4-tetrahydroquinolin-7-ol (12):** Compound **12** was prepared following **General protocol D** using compound **10** (50.0 mg, 0.228 mmol). Compound **12** was obtained (30.2 mg, 65%) was an amber oil. m/z: $(M+H)^+$ calculated for $C_{13}H_{20}NO^+$ 206.15, found 206.1 (ESI).

**8-ethyl-4,11,11-trimethyl-2,3,8,9,10,11-hexahydro-[1,4]oxazino[2,3-*b*]pyrido[2,3-*i*]phenoxazin-4-ium (25-64):** Compound **25-64** was prepared following **General protocol E** using compounds **12** (25.0 mg, 0.122 mmol) and **13** (40.0 mg, 0.122 mmol). Compound **25-64** was obtained (17.5 mg, 35%) as a blue solid. m/z: $(M+H)^+$ calculated for $C_{22}H_{26}N_3O_2^+$ 364.20, found 364.1 (ESI).

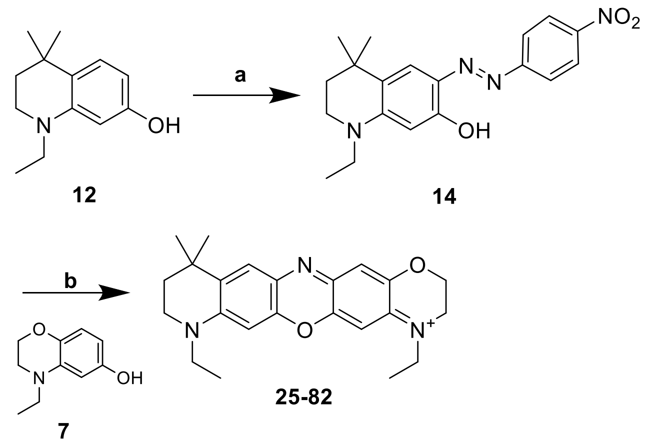


Scheme 4: Synthetic route to **25-82**. Reagents and conditions: a) (i) 2M hydrochloric acid / methanol (5:1), p-nitrobenzenediazonium tetrafluoroborate, 0 °C; (ii) $K_2CO_3$, 0 °C; b) acetic acid, 110 °C

**(E)-1-ethyl-4,4-dimethyl-6-((4-nitrophenyl)diazenyl)-1,2,3,4-tetrahydroquinolin-7-ol (14):** Compound **14** was prepared following **General protocol C** using compound **12** (30 mg, 0.146

mmol). Compound **14** was obtained as a red solid, which was used for the next step without further purification.

**4,8-diethyl-11,11-dimethyl-2,3,8,9,10,11-hexahydro-[1,4]oxazino[2,3-*b*]pyrido[2,3-*i*]phenoxazin-4-ium (25-82):** Compound **25-82** was prepared following **General protocol E** using compounds **14** (50.0 mg, 0.141mmol) and **7** (32.8 mg, 0.183mmol). Compound **25-82** was obtained (11.2 mg, 19%) as a blue solid. m/z: $(M+H)^+$ calculated for $C_{23}H_{28}N_3O_2^+$ 378.22, found 378.1 (ESI).

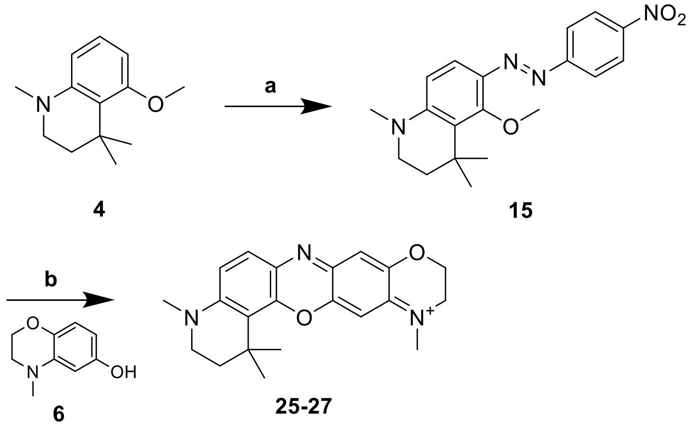


<u>**Scheme 5:**</u> Synthetic route to **25-27**. Reagents and conditions: a) (i) 2M hydrochloric acid / methanol (5:1), p-nitrobenzenediazonium tetrafluoroborate, 0 °C; (ii) $K_2CO_3$, 0 °C; b) acetic acid, 110 °C

**(E)-5-methoxy-1,4,4-trimethyl-6-((4-nitrophenyl)diazenyl)-1,2,3,4-tetrahydroquinoline (15):** Compound **15** was prepared following **General protocol C** using compound **4** (130.0 mg, 0.633 mmol). Compound **15** was obtained as a red solid, which was used for the next step without further purification.

**1,1,4,12-tetramethyl-1,2,3,4,10,11-hexahydro-[1,4]oxazino[2,3-*b*]pyrido[3,2-*h*]phenoxazin-12-ium (25-27):** Compound **25-27** was prepared following **General protocol E** using compounds **15** (55.7 mg, 0.151 mmol) and **6** (25.0 mg, 0.151 mmol). Compound **25-27** was obtained (19.1 mg, 45%) as a blue solid. m/z: $(M+H)^+$ calculated for $C_{21}H_{24}N_3O_2^+$ 350.19, found 350.1 (ESI).

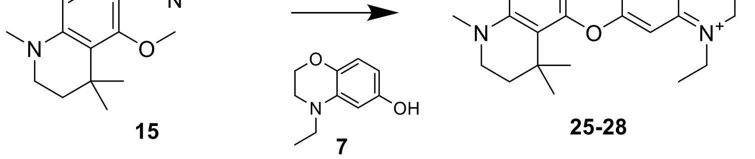


<u>**Scheme 6:**</u> Synthetic route to **25-28**. Reagents and conditions: a) acetic acid, 110 °C.

**12-ethyl-1,1,4-trimethyl-1,2,3,4,10,11-hexahydro-[1,4]oxazino[2,3-*b*]pyrido[3,2-*h*]phenoxazin-12-ium (25-28):** Compound **25-28** was prepared following **General protocol E** using compounds **15** (24.7 mg, 0.070 mmol) and **7** (12.5 mg, 0.070 mmol). Compound **25-28** was obtained (15.0 mg, 52%) as a blue solid. m/z: $(M+H)^+$ calculated for $C_{22}H_{26}N_3O_2^+$ 364.20, found 364.1 (ESI).

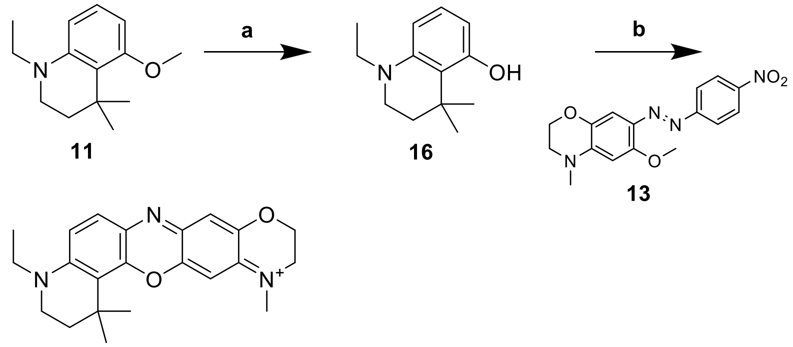


**Scheme 7:** Synthetic route to **25-88**. Reagents and conditions: a) 1 M boron tribromide, dichloromethane, 0 °C to rt; b) acetic acid, 110 °C.

**1-ethyl-4,4-dimethyl-1,2,3,4-tetrahydroquinolin-5-ol (16):** Compound **16** was prepared following **General protocol D** using compound **11** (50.0 mg, 0.228 mmol). Compound **16** was obtained (26.2 mg, 56%) as an orange oil. m/z: $(M+H)^+$ calculated for $C_{13}H_{20}NO^+$ 206.15, found 206.1 (ESI).

**4-ethyl-1,1,12-trimethyl-1,2,3,4,10,11-hexahydro-[1,4]oxazino[2,3-*b*]pyrido[3,2-*h*]phenoxazin-12-ium (25-88):** Compound **25-88** was prepared following **General protocol E** using compounds **16** (10.7 mg, 0.052 mmol) and **13** (13.6 mg, 0.040 mmol). Compound **25-88** was obtained (13.3 mg, 81%) as a blue solid. m/z: $(M+H)^+$ calculated for $C_{22}H_{26}N_3O_2^+$ 364.20, found 364.2 (ESI).

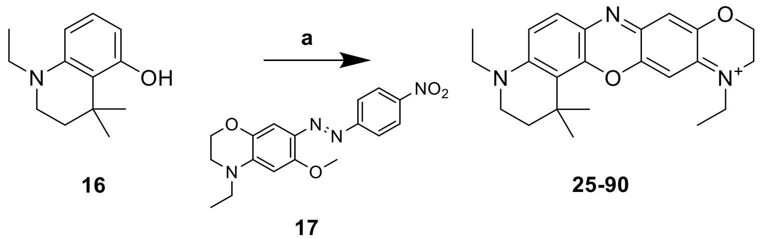


**Scheme 8:** Synthetic route to **25-90**. Reagents and conditions: a) acetic acid, 110 °C.

Compound **17** were synthesized following protocols reported by Wang[2].

**4,12-diethyl-1,1-dimethyl-1,2,3,4,10,11-hexahydro-[1,4]oxazino[2,3-*b*]pyrido[3,2-*h*]phenoxazin-12-ium (25-90):** Compound **25-90** was prepared following **General protocol E** using compounds **16** (10.9 mg, 0.053 mmol) and **17** (14.0 mg, 0.041 mmol). Compound **25-90**

was obtained (15.8 mg, 91%) as a blue solid. m/z: $(M+H)^+$ calculated for $C_{23}H_{28}N_3O_2^+$ 378.22, found 378.2 (ESI).

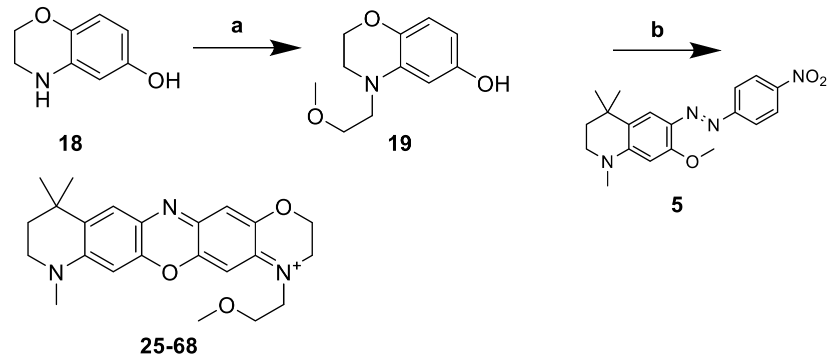

Scheme 9: Synthetic route to **25-68**. Reagents and conditions: a) (i) 2-methoxyacetyl chloride, acetonitrile 0 °C to rt, (ii) 1M borane-tetrahydrofuran, tetrahydrofuran 0 °C to rt; b) acetic acid, 110 °C.

Compound **26** were synthesized following protocols reported by Wang[2]

**4-(2-methoxyethyl)-3,4-dihydro-2H-benzo[*b*][1,4]oxazin-6-ol (19):** Compound **19** was prepared by suspending compound **18** (200.0 mg, 1.32 mmol) in 5 mL acetonitrile, 2-methoxyacetyl chloride (143.6 mg, 120.6 mmol) was then added dropwise at 0 °C and slowly warmed to rt. The reaction mixture was stirred overnight. The solvent was removed under a vacuum, and the intermediate was suspended in 5.0 mL of tetrahydrofuran and 5.0 mL of 1M borane-tetrahydrofuran was added dropwise to the solution at 0 °C and stirred overnight and slowly warmed to rt. The solution was then placed in a 0 °C bath and the borane-tetrahydrofuran was destroyed with methanol being added dropwise. The solvent was removed under a vacuum and was purified with flash column chromatography with silica gel. The residue was purified with a Ethyl Acetate/Hexane as the eluent to obtain compound (46.1 mg, 17%) a colorless oil. m/z: $(M+H)^+$ calculated for $C_{11}H_{16}NO_3^+$ 210.11, found 210.0 (ESI).

**4-(2-methoxyethyl)-8,11,11-trimethyl-2,3,8,9,10,11-hexahydro-[1,4]oxazino[2,3-*b*]pyrido[2,3-*i*]phenoxazin-4-ium (25-68):** Compound **25-68** was prepared following **General protocol E** using compounds **19** (25.0 mg, 0.119 mmol) and **5** (42.3 mg, 0.119 mmol). Compound **25-68** was obtained (19.5 mg, 37%) as a blue solid. m/z: $(M+H)^+$ calculated for $C_{23}H_{28}N_3O_3^+$ 394.21, found 394.1 (ESI).

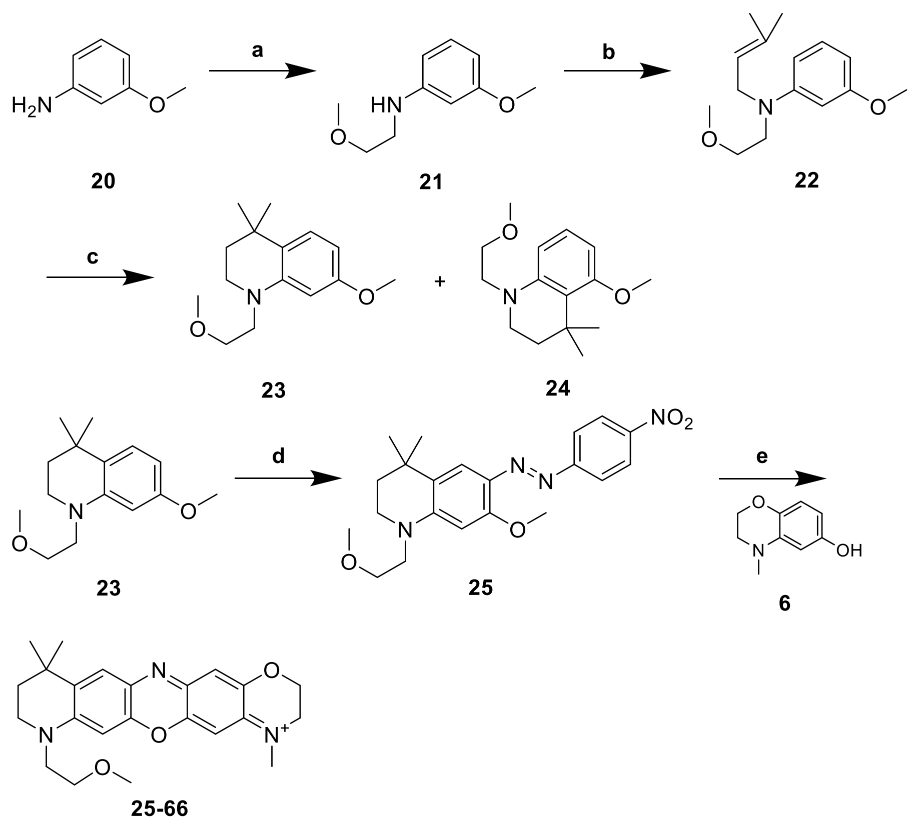

**Scheme 10:** Synthetic route to **25-66**. Reagents and conditions: a) (i) 2-methoxyacetyl chloride, acetonitrile 0 °C to rt, (ii) 1M borane-tetrahydrofuran, tetrahydrofuran 0 °C to rt; b) 1-bromo-3-methylbut-2-ene, $K_2CO_3$, acetonitrile, 80 °C; c) methanesulfonic acid, 80 °C; d) (i) 2M hydrochloric acid / methanol (5:1), p-nitrobenzenediazonium tetrafluoroborate, 0 °C; (ii) $K_2CO_3$, 0 °C; e) acetic acid, 110 °C.

**3-methoxy-N-(2-methoxyethyl)aniline (21):** Compound **21** was prepared by suspending compound **20** (5g, 40.60 mmol) in 60 mL acetonitrile, 2-methoxyacetyl chloride (4.41 g, 40.60 mmol) was then added dropwise at 0 °C and slowly warmed to rt. The reaction mixture was stirred overnight. The solvent was removed under a vacuum, and an aliquot of crude intermediate (3.50 g, 17.93 mmol) was suspended in 53.8 mL of tetrahydrofuran and 53.8 mL of 1M borane-tetrahydrofuran was added dropwise to the solution at 0 °C and stirred overnight and slowly warmed to rt. The solution was then placed in a 0 °C bath and the borane-tetrahydrofuran was destroyed with methanol being added. The solvent was removed under a vacuum and was purified with flash column chromatography with silica gel. The residue was purified with an Ethyl Acetate/Hexane as the eluent to obtain compound (3.03 g, 93%) an amber oil. m/z: $(M+H)^+$ calculated for $C_{10}H_{16}NO_2^+$ 182.12, found 182.0 (ESI).

**3-methoxy-N-(2-methoxyethyl)-N-(3-methylbut-2-en-1-yl)aniline (22):** Compound **22** was prepared following **General protocol A** using compound **21** (1.5 g, 8.28 mmol). Compound **22**

was obtained (945 mg, 46%) as a yellow oil. m/z: $(M+H)^+$ calculated for $C_{15}H_{24}NO_2^+$ 250.18, found 250.1 (ESI).

**7-methoxy-1-(2-methoxyethyl)-4,4-dimethyl-1,2,3,4-tetrahydroquinoline (23):** Compound **23** was prepared following **General protocol B** using compound **22** (500.0 mg, 2.01 mmol). Compound **23** was obtained (271.4 mg, 54%) as an amber oil. m/z: $(M+H)^+$ calculated for $C_{15}H_{24}NO_2^+$ 250.18, found 250.1 (ESI).

**5-methoxy-1-(2-methoxyethyl)-4,4-dimethyl-1,2,3,4-tetrahydroquinoline (24):** Compound **24** was prepared following **General protocol B** using compound **22** (500.0 mg, 2.01 mmol). Compound **24** was obtained (104.3 mg, 21%) as an amber oil m/z: $(M+H)^+$ calculated for $C_{15}H_{24}NO_2^+$ 250.18, found 250.1 (ESI).

**(E)-7-methoxy-1-(2-methoxyethyl)-4,4-dimethyl-6-((4-nitrophenyl)diazenyl)-1,2,3,4-tetrahydroquinoline (25):** Compound **25** was prepared following **General protocol C** using compound **23** (250.0 mg, 1.22 mmol). Compound **25** was obtained as a red solid, which was used for the next step without further purification.

**8-(2-methoxyethyl)-4,11,11-trimethyl-2,3,8,9,10,11-hexahydro-[1,4]oxazino[2,3-*b*]pyrido[2,3-*i*]phenoxazin-4-ium (25-66):** Compound **25-66** was prepared following **General protocol E** using compounds **25** (60.3 mg, 0.151 mmol) and **6** (25.0 mg, 0.151 mmol). Compound **25-66** was obtained (23.1 mg, 35%) as a blue solid. m/z: $(M+H)^+$ calculated for $C_{23}H_{28}N_3O_3^+$ 394.21, found 394.2 (ESI).

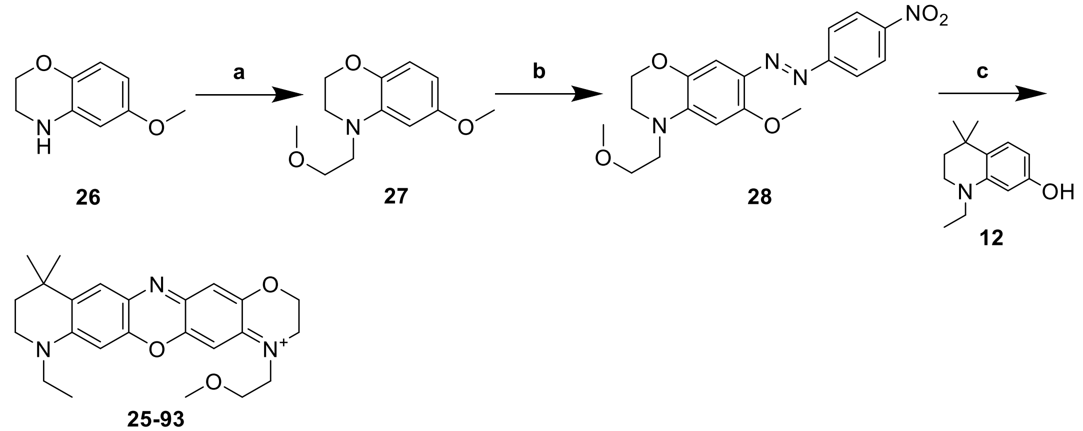


<u>**Scheme 11:**</u> Synthetic route to **25-93**. Reagents and conditions: a) (i) 2-methoxyacetyl chloride, acetonitrile 0 °C to rt, (ii) 1M borane-tetrahydrofuran, tetrahydrofuran 0 °C to rt; b) (i) 2M hydrochloric acid / methanol (5:1), p-nitrobenzenediazonium tetrafluoroborate, 0 °C; (ii) $K_2CO_3$, 0 °C; c) acetic acid, 110 °C.

Compound **26** were synthesized following protocols reported by Montaño[1].

**6-methoxy-4-(2-methoxyethyl)-3,4-dihydro-2H-benzo[*b*][1,4]oxazine (27):** Compound **27** was prepared by suspending compound **26** (90.0 mg, 0.544 mmol) in 5 mL acetonitrile, 2-

methoxyacetyl chloride (65.0 mg, 0.599 mmol) was then added dropwise at 0 °C and slowly warmed to rt. The reaction mixture was stirred overnight. The solvent was removed under a vacuum, and the intermediate was suspended in 1.63 mL of tetrahydrofuran and 1.63 mL of 1M borane-tetrahydrofuran was added dropwise to the solution at 0 °C and stirred overnight and slowly warmed to rt. The solution was then placed in a 0 °C bath and the borane-tetrahydrofuran was destroyed with methanol being added. The solvent was removed under a vacuum and was purified with flash column chromatography with silica gel. The residue was purified with an Ethyl Acetate/Hexane as the eluent to obtain compound **26** (102.3 mg, 84%) an amber oil. m/z: $(M+H)^+$ calculated for $C_{12}H_{18}NO_3^+$ 224.13, found 224.0 (ESI).

**(E)-6-methoxy-4-(2-methoxyethyl)-7-((4-nitrophenyl)diazenyl)-3,4-dihydro-2H-benzo[*b*][1,4]oxazine (28):** Compound **28** was prepared following **General protocol C** using compound **27** (500.0 mg, 2.23 mmol). Compound **28** was obtained as a dark red solid, which was used for the next step without further purification.

**8-ethyl-4-(2-methoxyethyl)-11,11-dimethyl-2,3,8,9,10,11-hexahydro-[1,4]oxazino[2,3-*b*]pyrido[2,3-*i*]phenoxazin-4-ium (25-93):** Compound **25-93** was prepared following **General protocol E** using compounds **28** (21.5 mg, 0.057 mmol) and **12** (15.4 mg, 0.0 mmol). Compound **25-93** was obtained (14.6 mg, 56%) as a blue solid. m/z: $(M+H)^+$ calculated for $C_{24}H_{30}N_3O_3^+$ 408.23, found 408.2 (ESI).

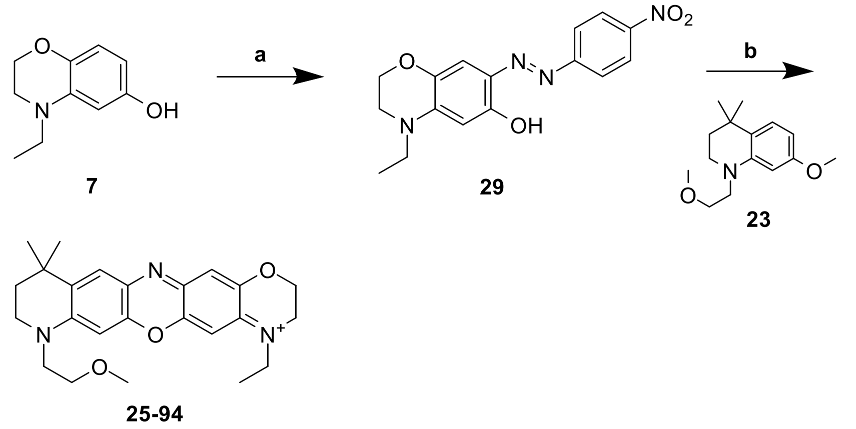

**Scheme 12:** Synthetic route to **25-94**. Reagents and conditions: a) (i) 2M hydrochloric acid / methanol (5:1), p-nitrobenzenediazonium tetrafluoroborate, 0 °C; (ii) $K_2CO_3$, 0 °C; b) acetic acid, 110 °C.

**(E)-4-ethyl-7-((4-nitrophenyl)diazenyl)-3,4-dihydro-2H-benzo[*b*][1,4]oxazin-6-ol (29):** Compound **29** was prepared following **General protocol C** using compound **7** (25.0 mg, 0.140 mmol). Compound **29** was obtained as a dark red solid, which was used for the next step without further purification.

**4-ethyl-8-(2-methoxyethyl)-11,11-dimethyl-2,3,8,9,10,11-hexahydro-[1,4]oxazino[2,3-*b*]pyrido[2,3-*i*]phenoxazin-4-ium (25-94):** Compound **25-94** was prepared following **General**

**protocol E** using compounds **29** (16.2 mg, 0.049 mmol) and **23** (16.0 mg, 0.064 mmol). Compound **25-94** was obtained (5.1 mg, 23%) as a blue solid. m/z: $(M+H)^+$ calculated for $C_{24}H_{30}N_3O_3^+$ 408.23, found 408.2 (ESI).

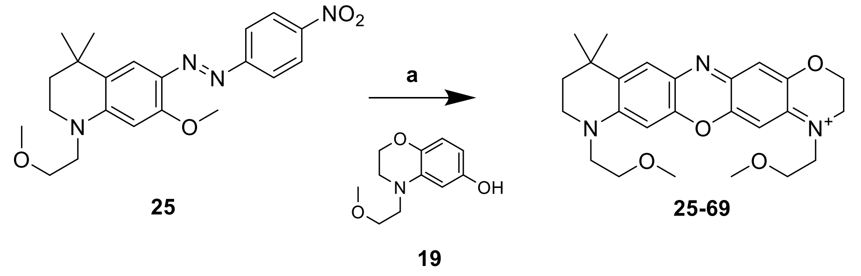

**Scheme 13:** Synthetic route to **25-69**. Reagents and conditions: a) acetic acid, 110 °C.

**4,8-bis(2-methoxyethyl)-11,11-dimethyl-2,3,8,9,10,11-hexahydro-[1,4]oxazino[2,3-*b*]pyrido[2,3-*i*]phenoxazin-4-ium (25-69):** Compound **25-69** was prepared following **General protocol E** using compounds **25** (47.6 mg, 0.119 mmol) and **19** (25.0 mg, 0.119 mmol). Compound **25-69** was obtained (43.4 mg, 75%) as a blue solid. m/z: $(M+H)^+$ calculated for $C_{25}H_{32}N_3O_4^+$ 438.24, found 438.2 (ESI).

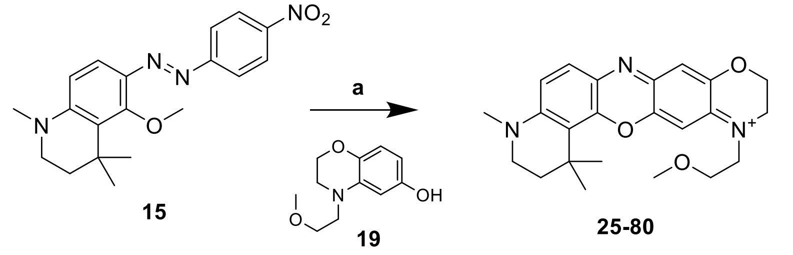

**Scheme 14:** Synthetic route to **25-80**. Reagents and conditions: a) acetic acid, 110 °C.

**12-(2-methoxyethyl)-1,1,4-trimethyl-1,2,3,4,10,11-hexahydro-[1,4]oxazino[2,3-*b*]pyrido[3,2-*h*]phenoxazin-12-ium (25-80):** Compound **25-80** was prepared following **General protocol E** using compounds **15** (15.0 mg, 0.042 mmol) and **19** (11.5 mg, 0.055 mmol). Compound **25-80** was obtained (7.2 mg, 38.7%) as a blue solid. m/z: $(M+H)^+$ calculated for $C_{23}H_{28}N_3O_3^+$ 394.21, found 394.2 (ESI).

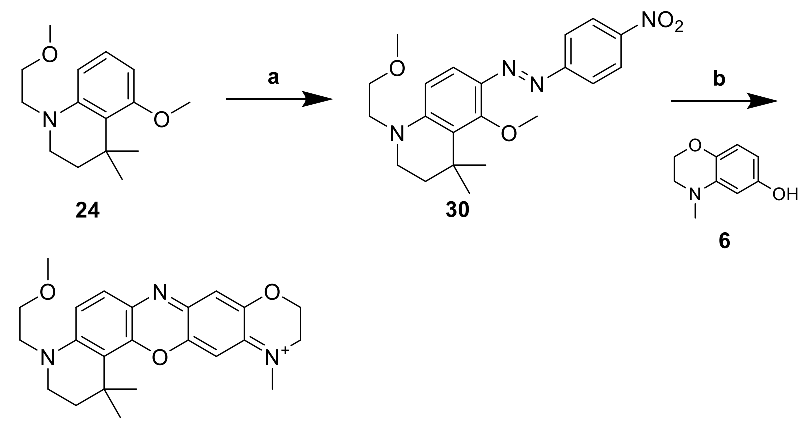


24 30 6

26-10

<u>Scheme 15:</u> Synthetic route to **26-10**. Reagents and conditions: a) (i) 2M hydrochloric acid / methanol (5:1), p-nitrobenzenediazonium tetrafluoroborate, 0 °C; (ii) $K_2CO_3$, 0 °C; b) acetic acid, 110 °C.

**(E)-5-methoxy-1-(2-methoxyethyl)-4,4-dimethyl-6-((4-nitrophenyl)diazenyl)-1,2,3,4-tetrahydroquinoline (30):** Compound **30** was prepared following **General protocol C** using compound **24** (50.0 mg, 0.201 mmol). Compound **30** was obtained as a red solid, which was used for the next step without further purification.

**4-(2-methoxyethyl)-1,1,12-trimethyl-1,2,3,4,10,11-hexahydro-[1,4]oxazino[2,3-*b*]pyrido[3,2-*h*]phenoxazin-12-ium (26-10):** Compound **26-10** was prepared following **General protocol E** using compounds **30** (16.4 mg, 0.041 mmol) and **6** (10.2 mg, 0.062 mmol). Compound **26-10** was obtained (5.9 mg, 33%) as a blue solid. . m/z: $(M+H)^+$ calculated for $C_{23}H_{28}N_3O_3^+$ 394.21, found 394.2 (ESI).

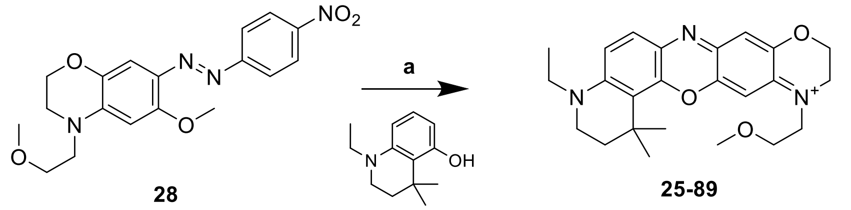


28 25-89

11

<u>Scheme 16:</u> Synthetic route to **25-89**. Reagents and conditions: a) acetic acid, 110 °C.

**4-ethyl-12-(2-methoxyethyl)-1,1-dimethyl-1,2,3,4,10,11-hexahydro-[1,4]oxazino[2,3-*b*]pyrido[3,2-*h*]phenoxazin-12-ium (25-89)** Compound **25-89** was prepared following **General protocol E** using compounds **28** (12.1 mg, 0.033 mmol) and **11** (8.7 mg, 0.042 mmol). Compound **25-89** was obtained (13.6 mg, 92%) as a blue solid. m/z: $(M+H)^+$ calculated for $C_{24}H_{30}N_3O_3^+$ 408.23, found 408.2 (ESI).

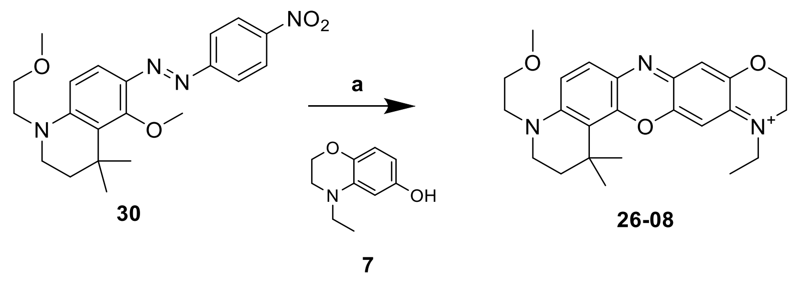

Scheme 17: Synthetic route to **26-08**. Reagents and conditions: a) acetic acid, 110 °C.

**12-ethyl-4-(2-methoxyethyl)-1,1-dimethyl-1,2,3,4,10,11-hexahydro-[1,4]oxazino[2,3-*b*]pyrido[3,2-*h*]phenoxazin-12-ium (26-08):** Compound **26-08** was prepared following **General protocol E** using compounds **30** (15.0 mg, 0.037 mmol) and **7** (8.8 mg, 0.049 mmol). Compound **26-08** was obtained (7.3 mg, 43%) as a blue solid. . m/z: $(M+H)^+$ calculated for $C_{24}H_{30}N_3O_3^+$ 408.23, found 408.2 (ESI).

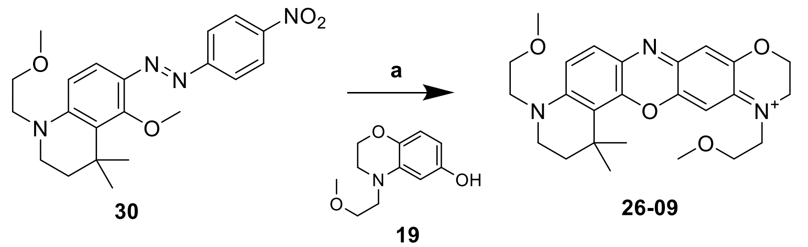

Scheme 18: Synthetic route to **26-09**. Reagents and conditions: a) acetic acid, 110 °C.

**4,12-bis(2-methoxyethyl)-1,1-dimethyl-1,2,3,4,10,11-hexahydro-[1,4]oxazino[2,3-*b*]pyrido[3,2-*h*]phenoxazin-12-ium (26-09):** Compound **26-09** was prepared following **General protocol E** using compounds **30** (15.0 mg, 0.037 mmol) and **19** (10.2 mg, 0.049 mmol). Compound **26-09** was obtained (6.3 mg, 35%) as a blue solid. m/z: $(M+H)^+$ calculated for $C_{25}H_{32}N_3O_4^+$ 438.24, found 438.2 (ESI).

2

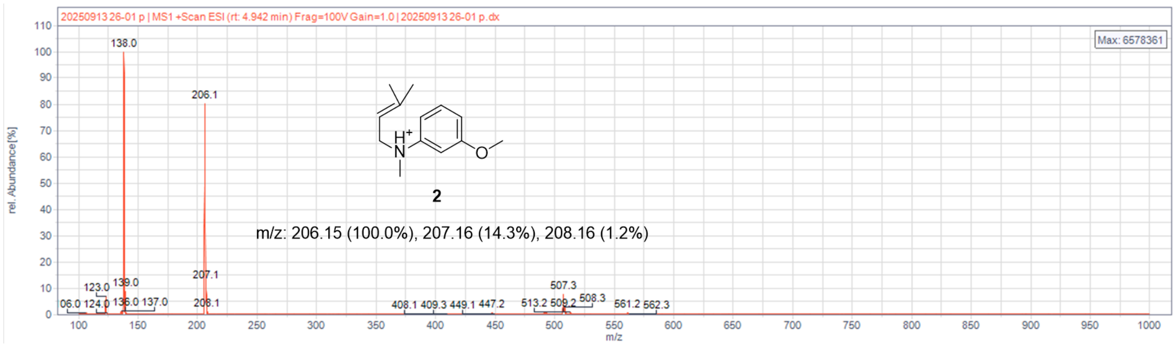


**Figure S1: HPLC-MS data for compound 2**

3

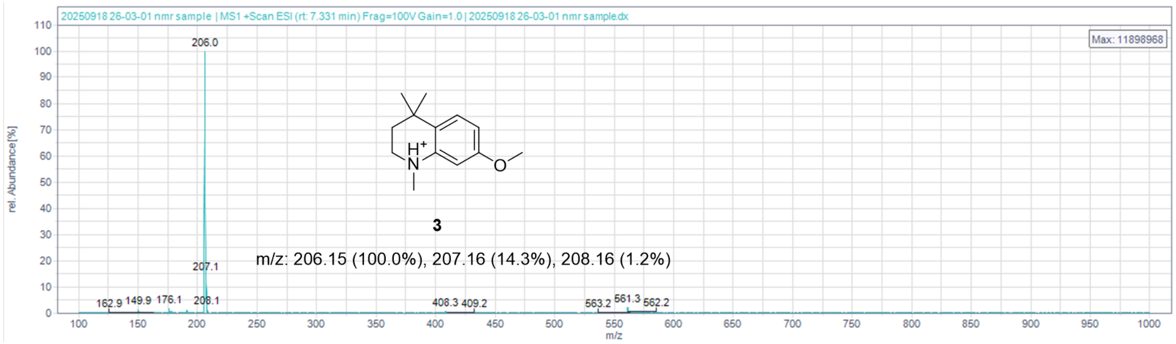


**Figure S2: HPLC-MS data for compound 3**

4

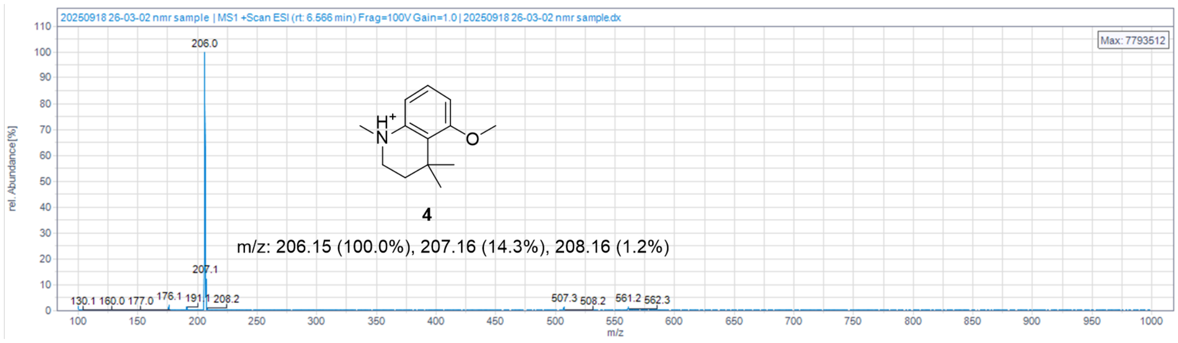


**Figure S3: HPLC-MS data for compound 4**

9

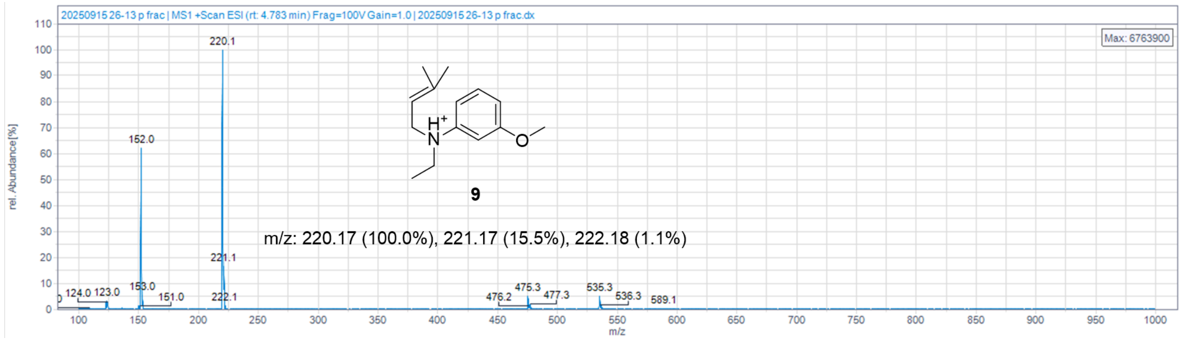


**Figure S4: HPLC-MS data for compound 9**

10

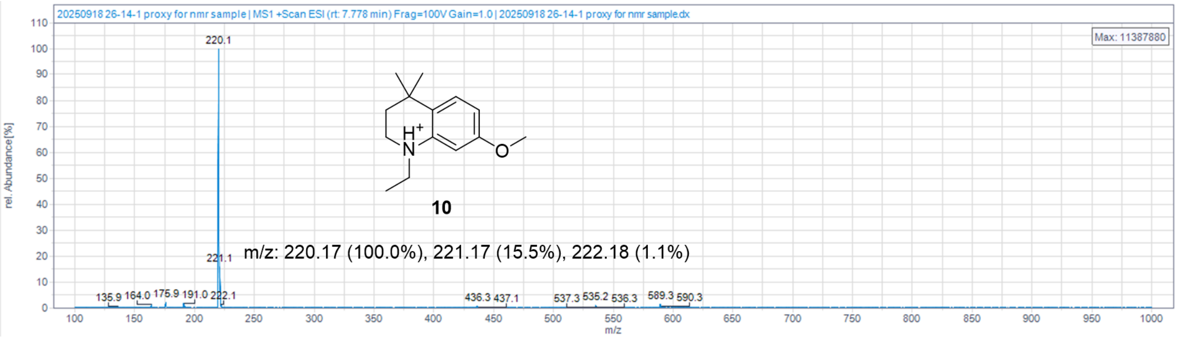


**Figure S5: HPLC-MS data for compound 10**

11

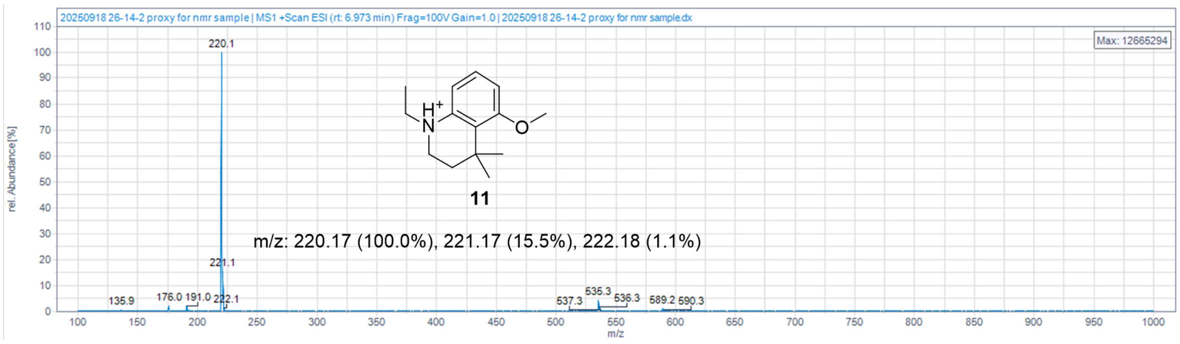


**Figure S6: HPLC-MS data for compound 11**

12

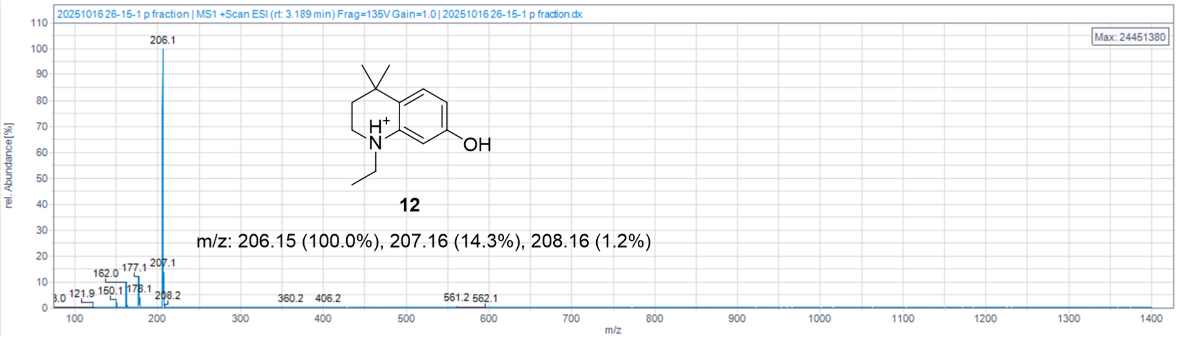


**Figure S7: HPLC-MS data for compound 12**

16

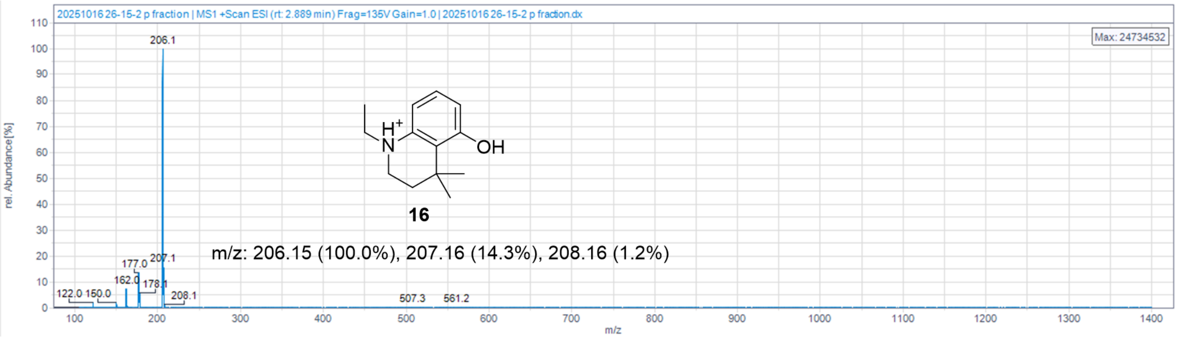


**Figure S8: HPLC-MS data for compound 16**

19

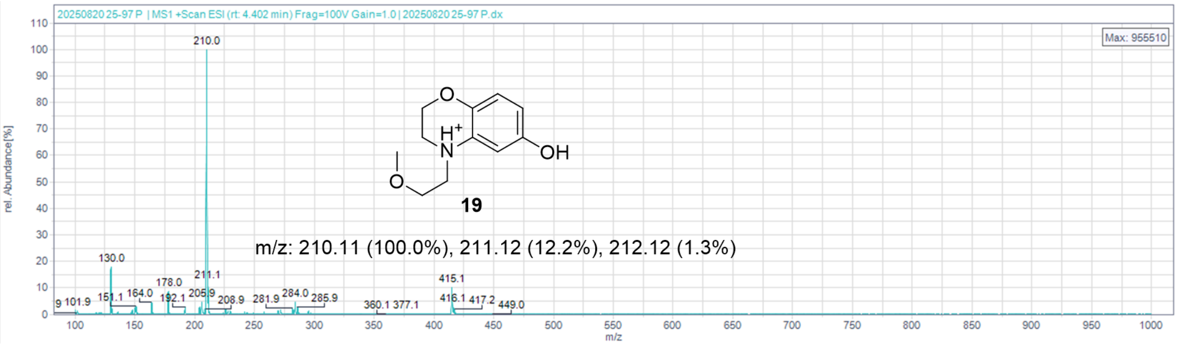


Figure S9: HPLC-MS data for compound 19

21

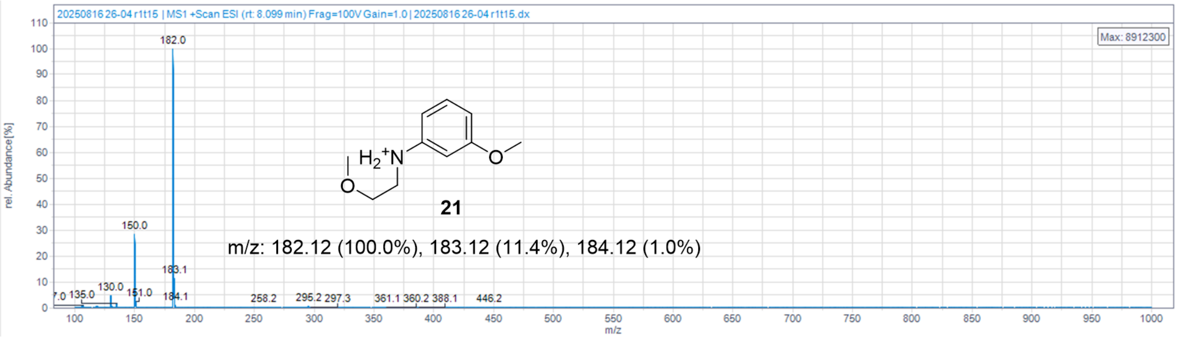


Figure S10: HPLC-MS data for compound 21

22

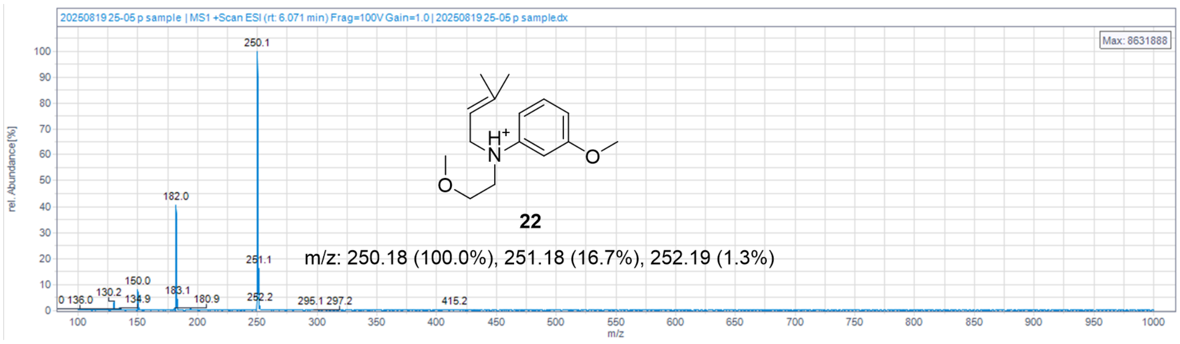


Figure S11: HPLC-MS data for compound 22

23

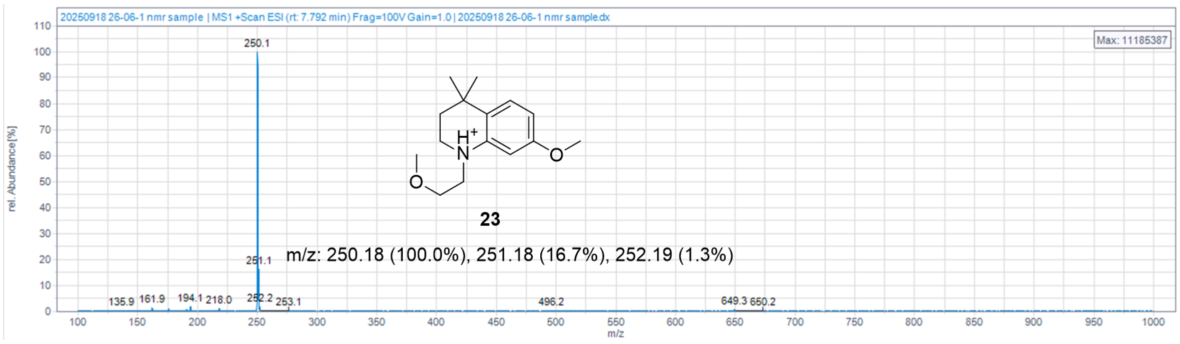


Figure S12: HPLC-MS data for compound 23

24

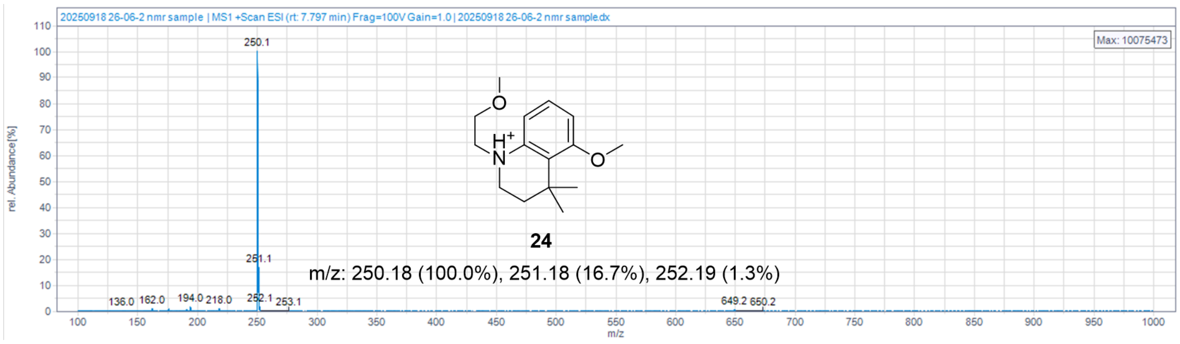


**Figure S13: HPLC-MS data for compound 24**

27

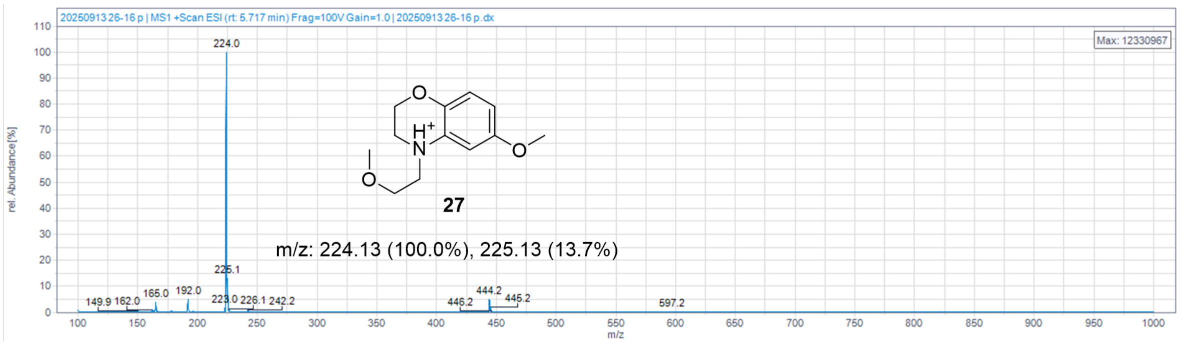


**Figure S14: HPLC-MS data for compound 27**

**Figure S1-14: HPLC-MS and purity analysis of oxazine dyes.** High pressure liquid chromatography-mass spectrometry (HPLC-MS) was used to identify mass of each compound.

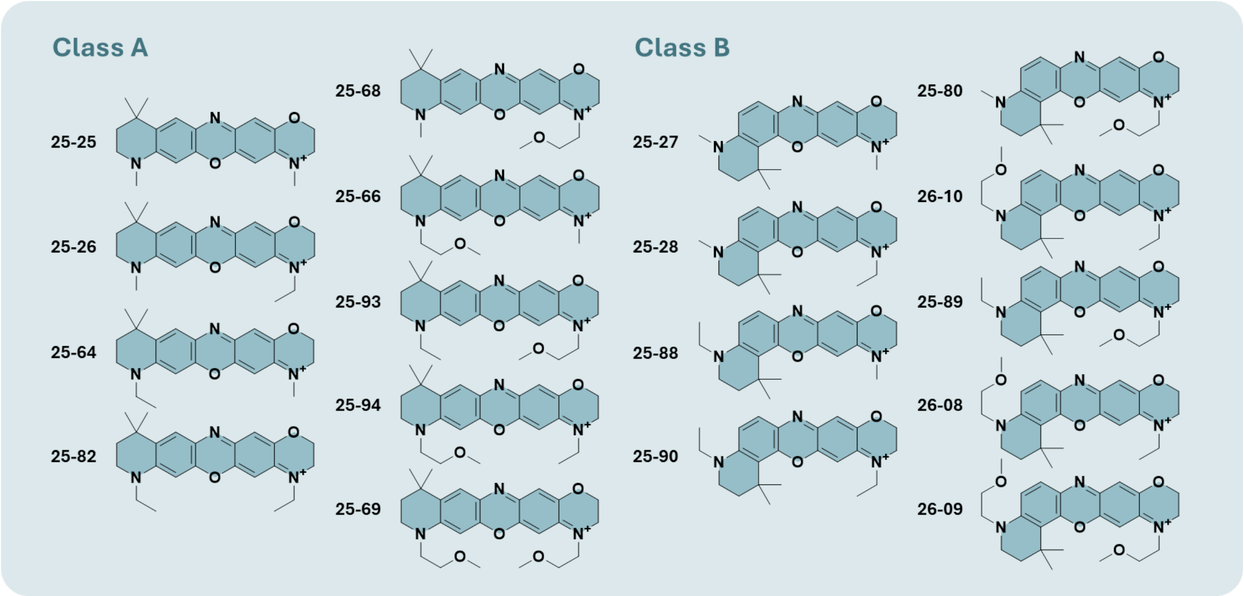


**Figure S15: Oxazine derivative library chemical structures.** Structures for 18 fluorophores in library, divided into two classes.

25-25

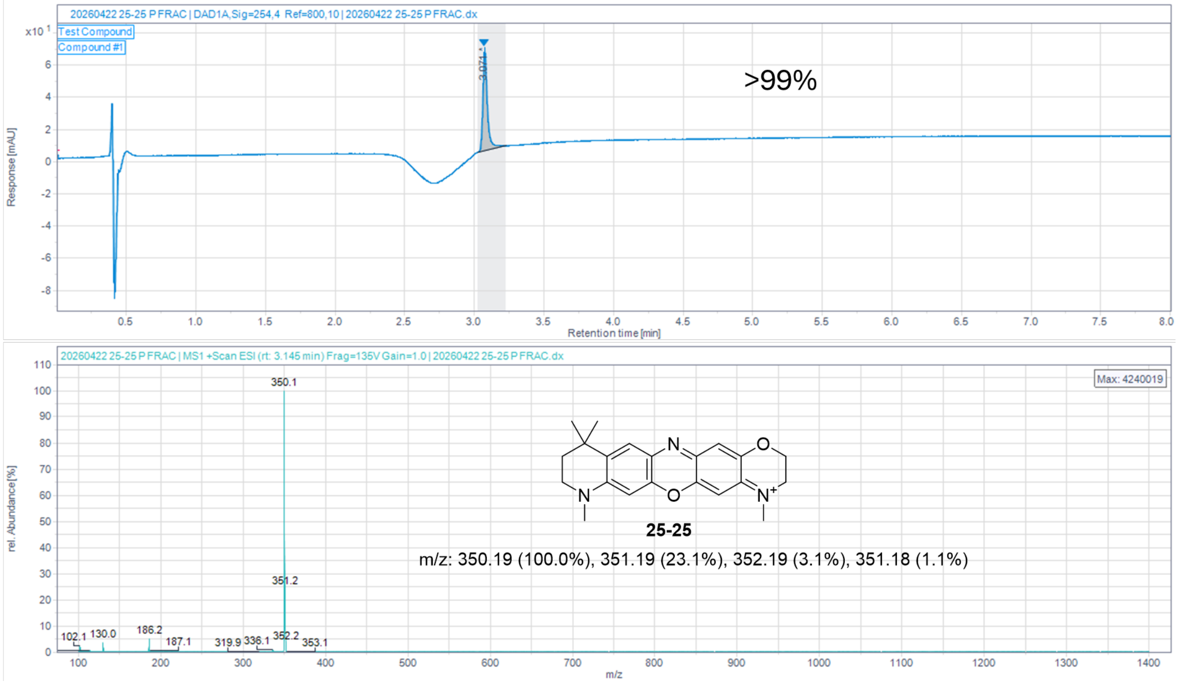


**Figure S16: HPLC-MS and purity analysis data for compound 25-25**

25-26

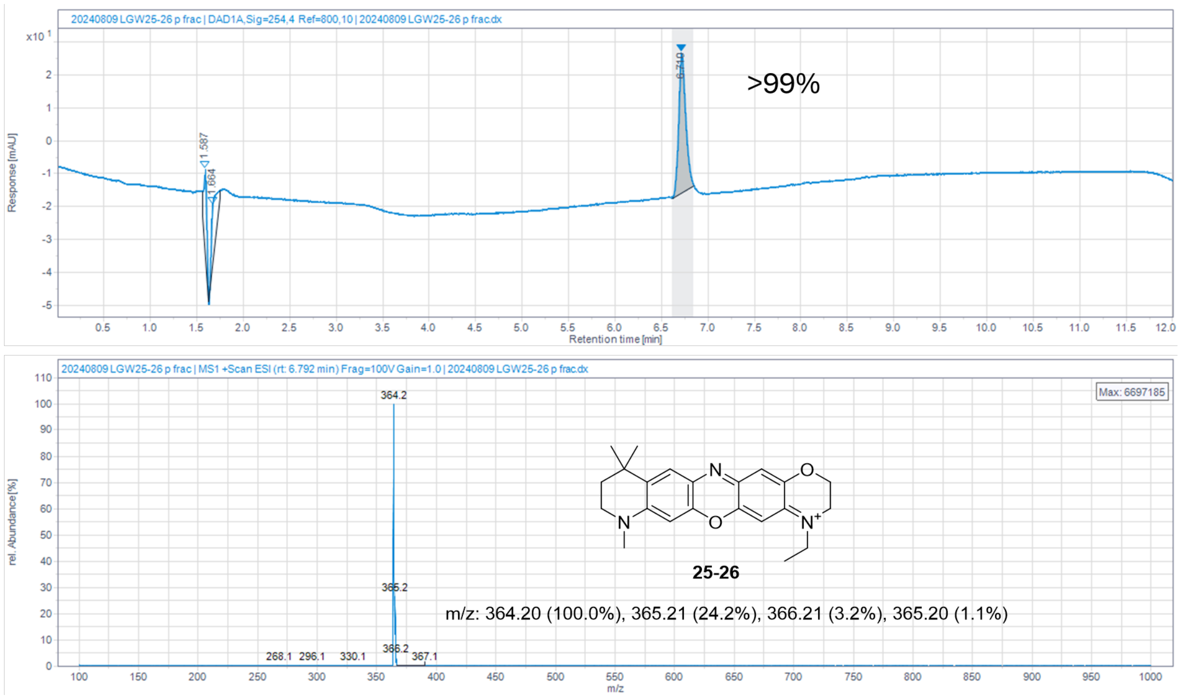


**Figure S17: HPLC-MS and purity analysis data for compound 25-26**

25-64

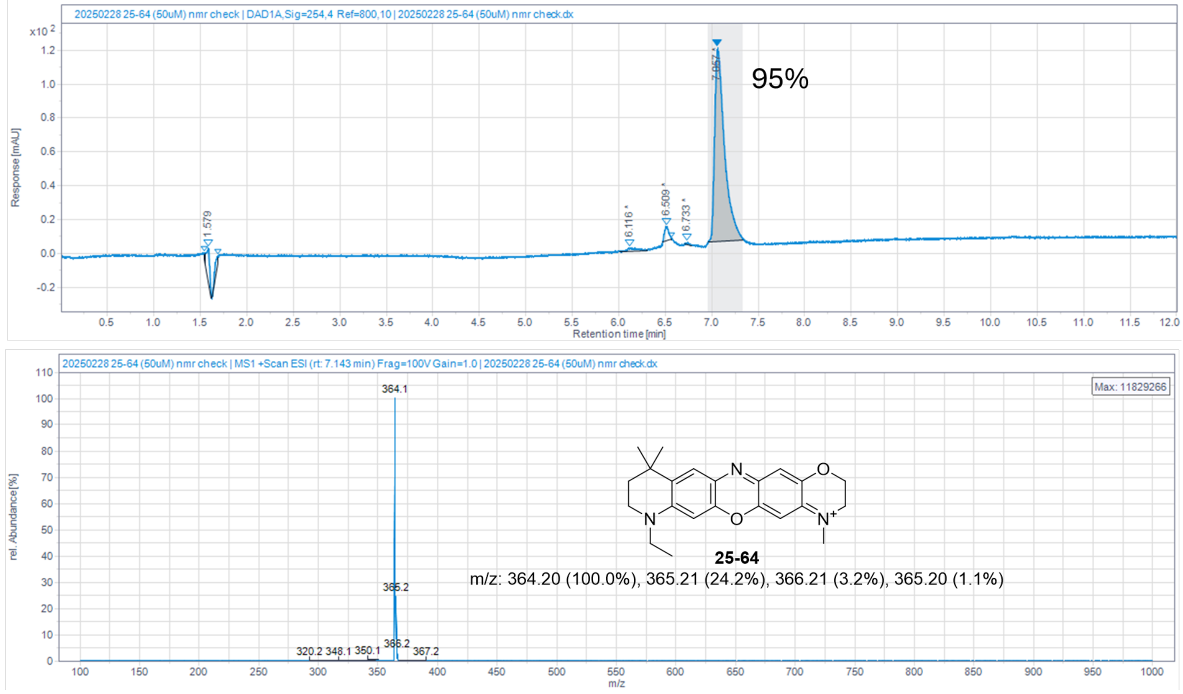


**Figure S18: HPLC-MS and purity analysis data for compound 25-64**

25-82

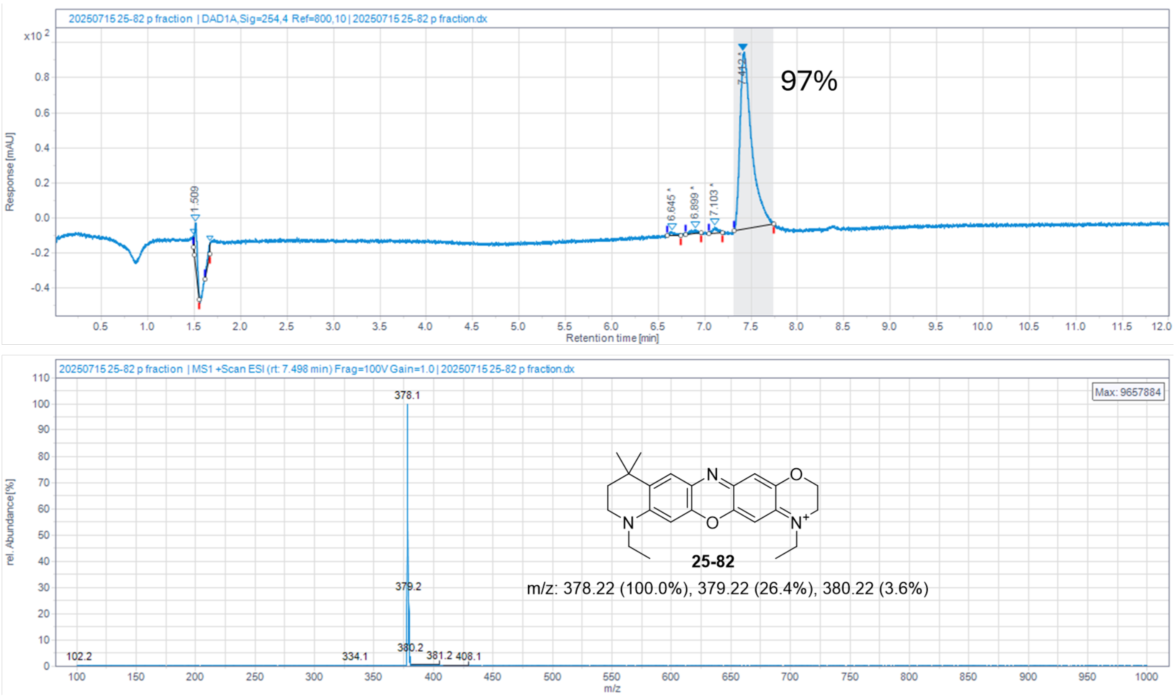


**Figure S19: HPLC-MS and purity analysis data for compound 25-82**

25-68

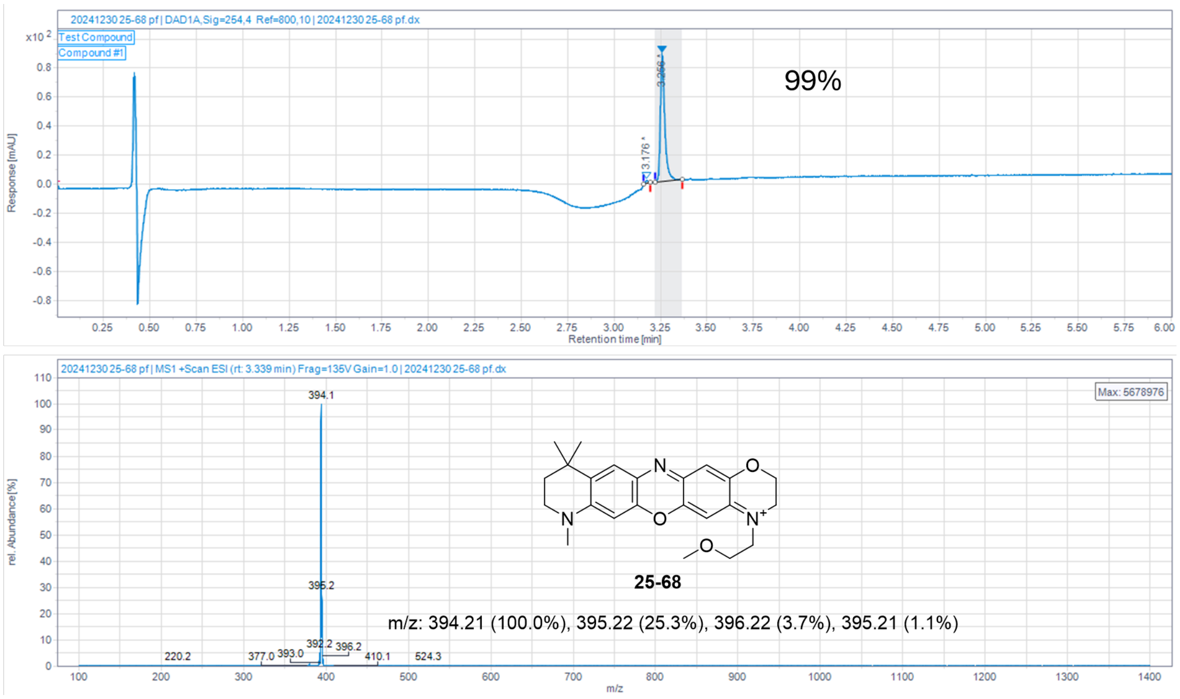


**Figure S20: HPLC-MS and purity analysis data for compound 25-68**

25-66

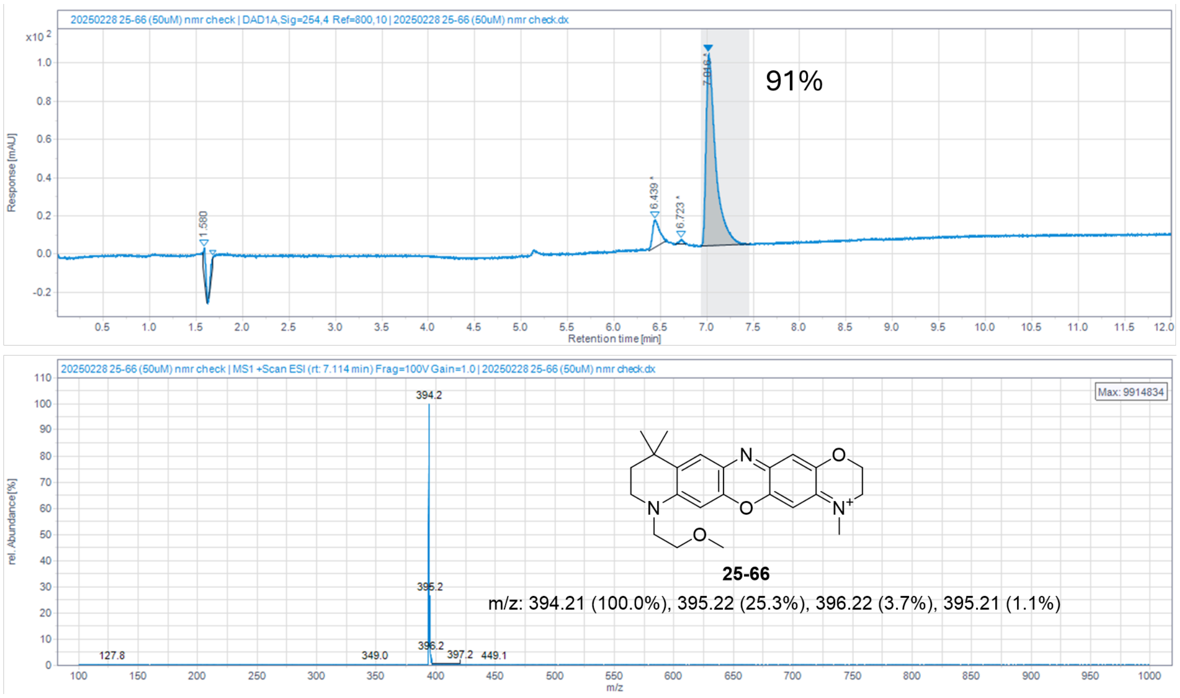


**Figure S21: HPLC-MS and purity analysis data for compound 25-66**

25-93

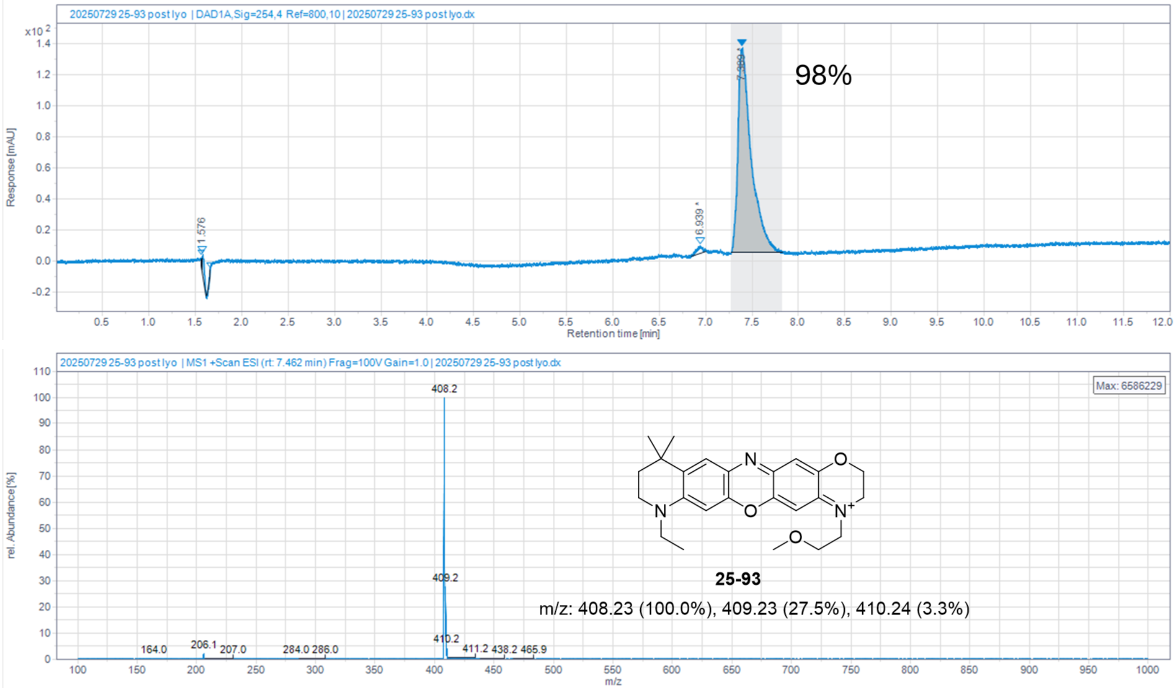


**Figure S22: HPLC-MS and purity analysis data for compound 25-93**

25-94

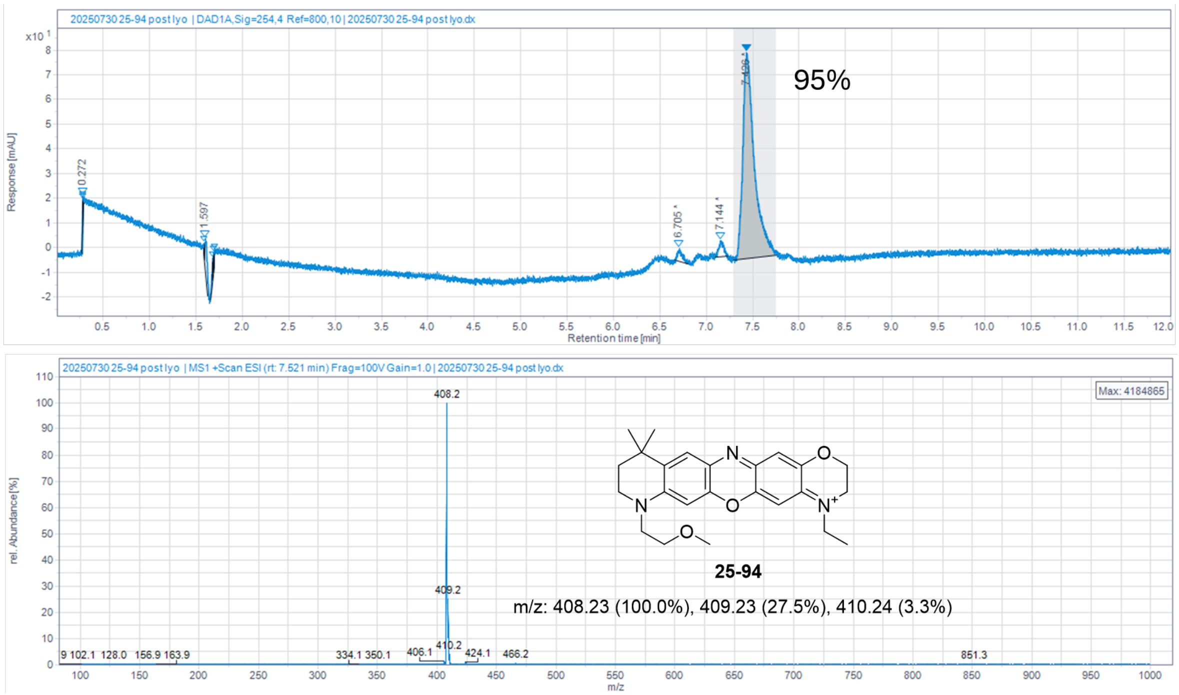


**Figure S23: HPLC-MS and purity analysis data for compound 25-94**

25-69

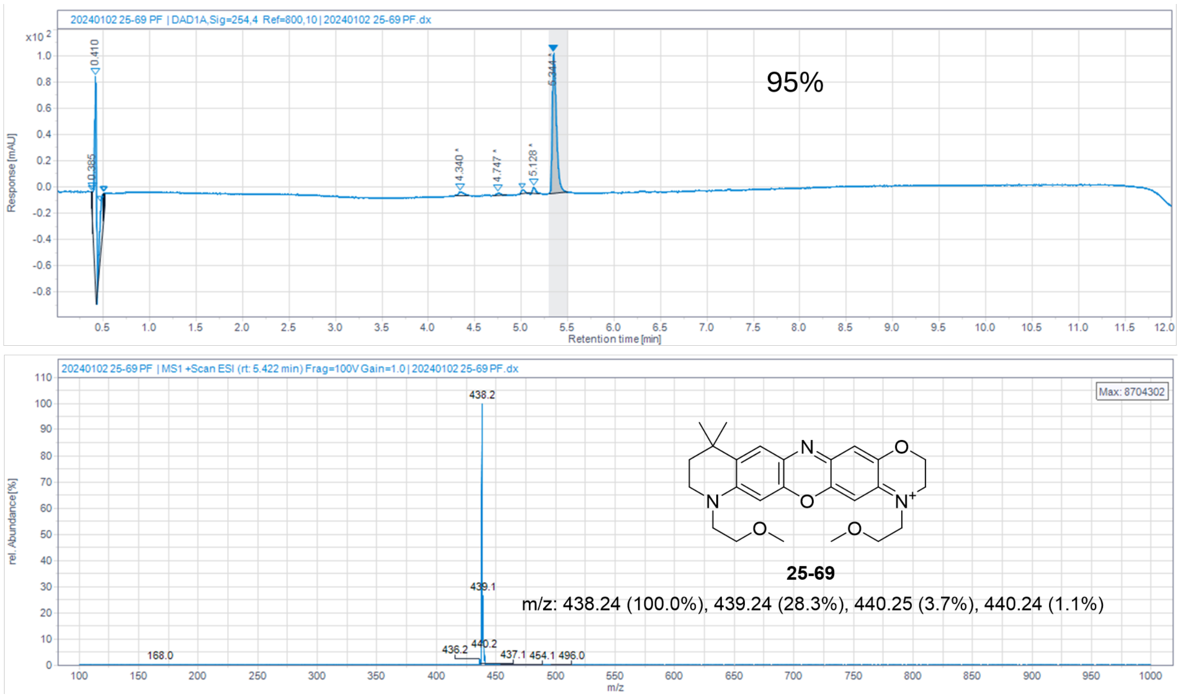


**Figure S24: HPLC-MS and purity analysis data for compound 25-69**

25-27

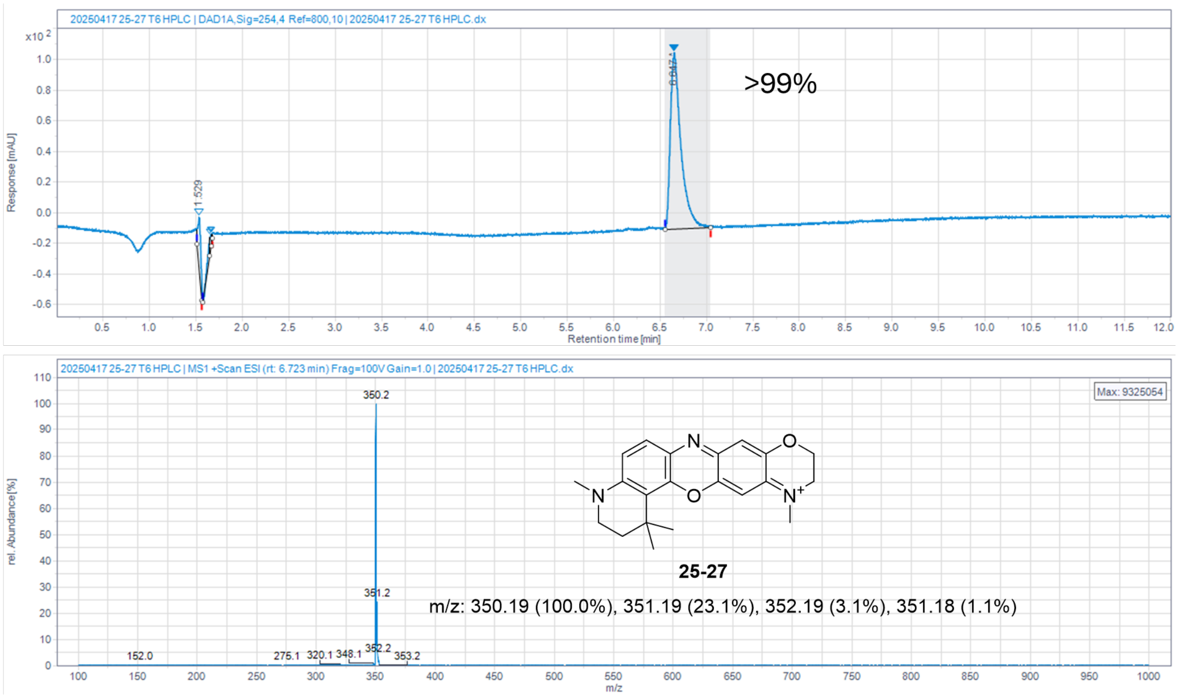


**Figure S25: HPLC-MS and purity analysis data for compound 25-27**

25-28

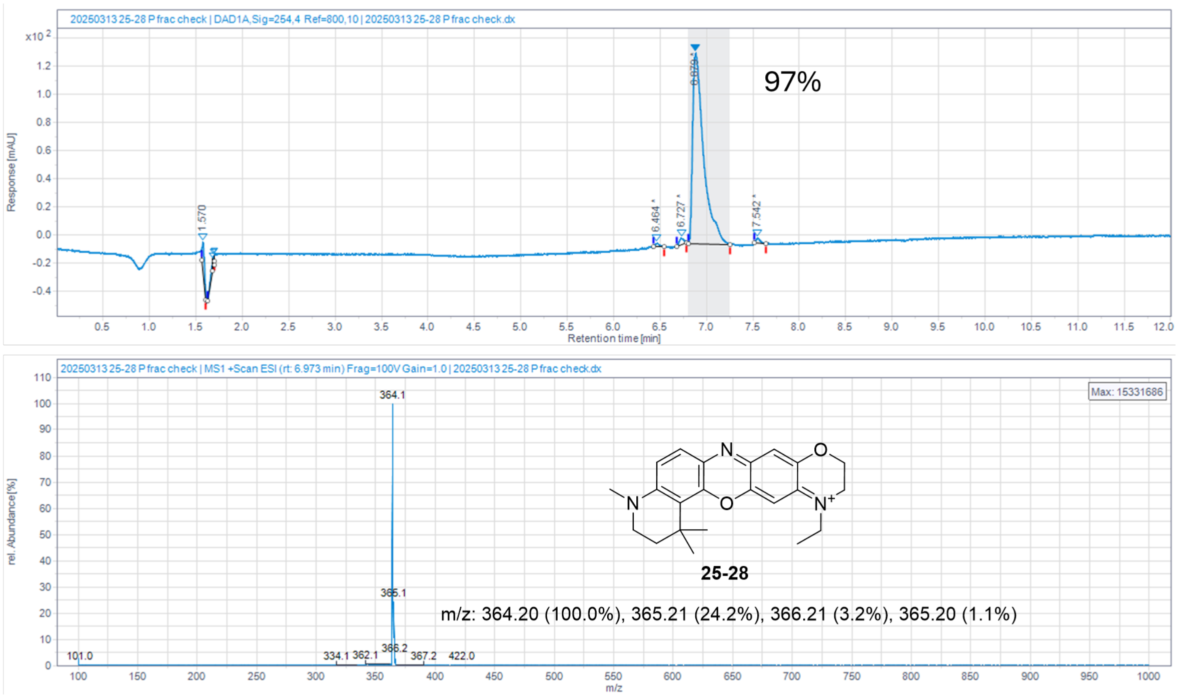


**Figure S26: HPLC-MS and purity analysis data for compound 25-28**

25-88

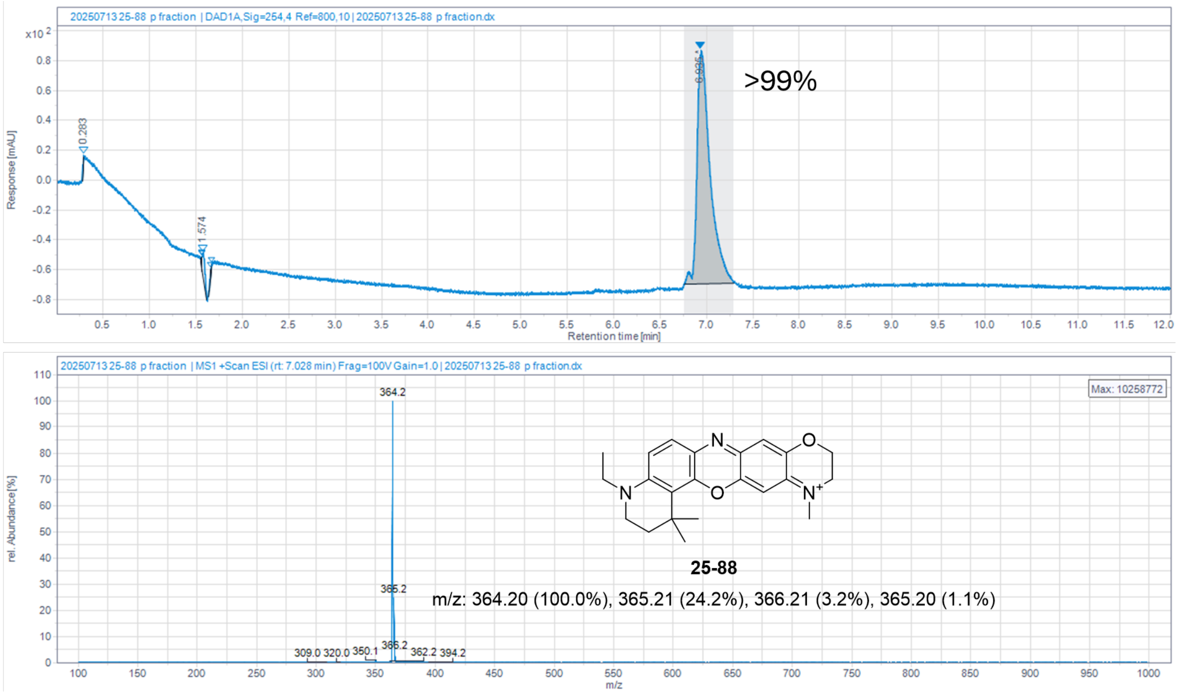


**Figure S27: HPLC-MS and purity analysis data for compound 25-88**

25-90

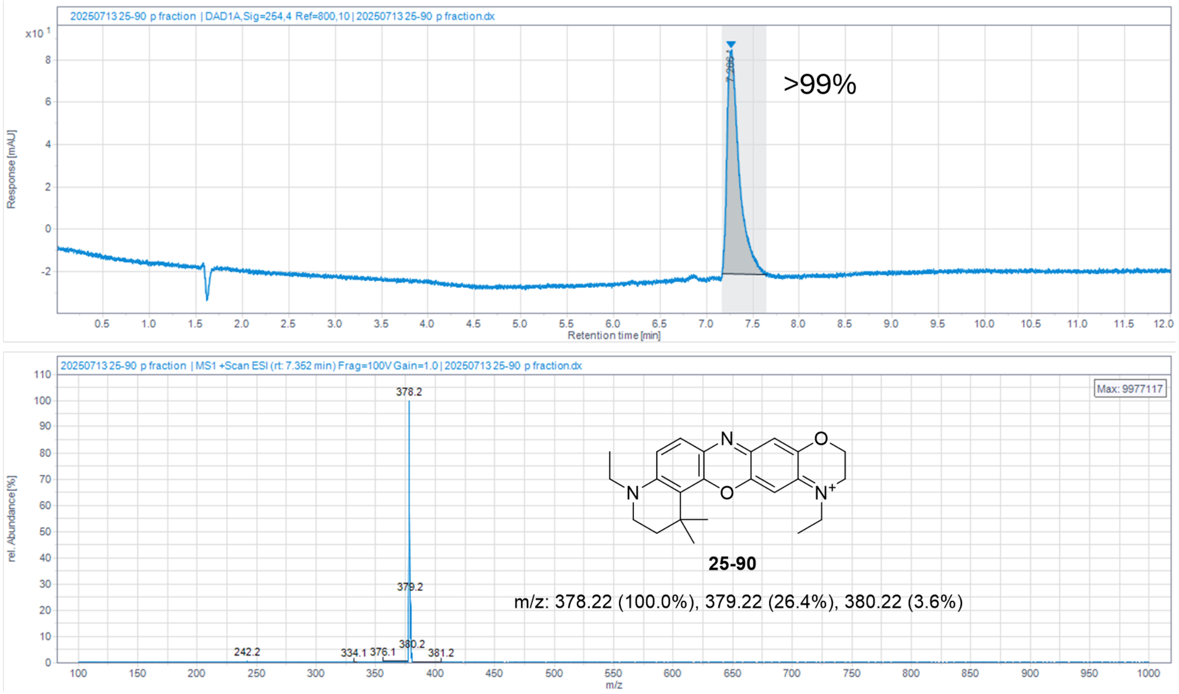


**Figure S28: HPLC-MS and purity analysis data for compound 25-90**

25-80

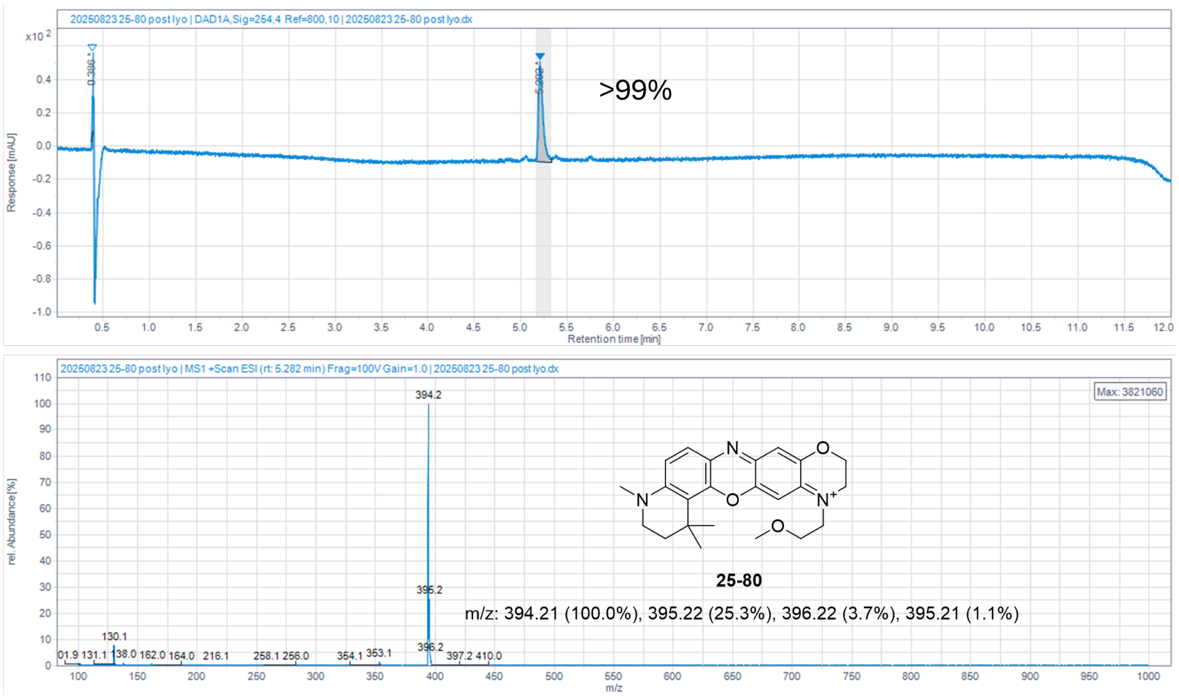


**Figure S29: HPLC-MS and purity analysis data for compound 25-80**

26-10

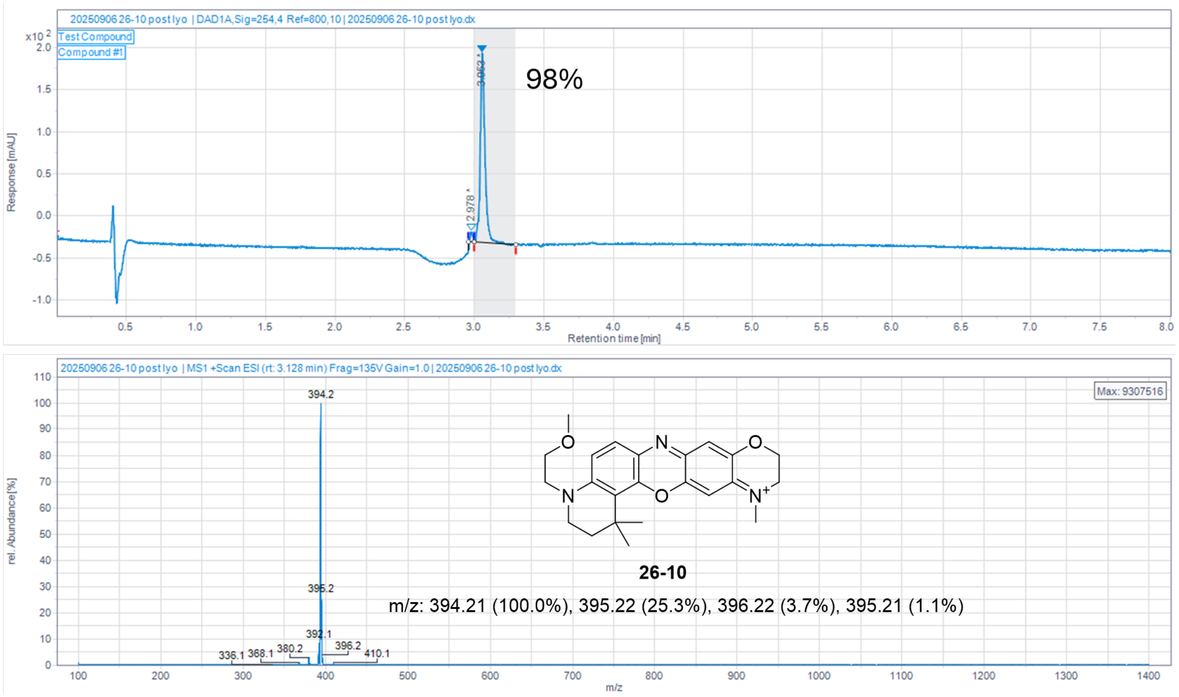


**Figure S30: HPLC-MS and purity analysis data for compound 26-10**

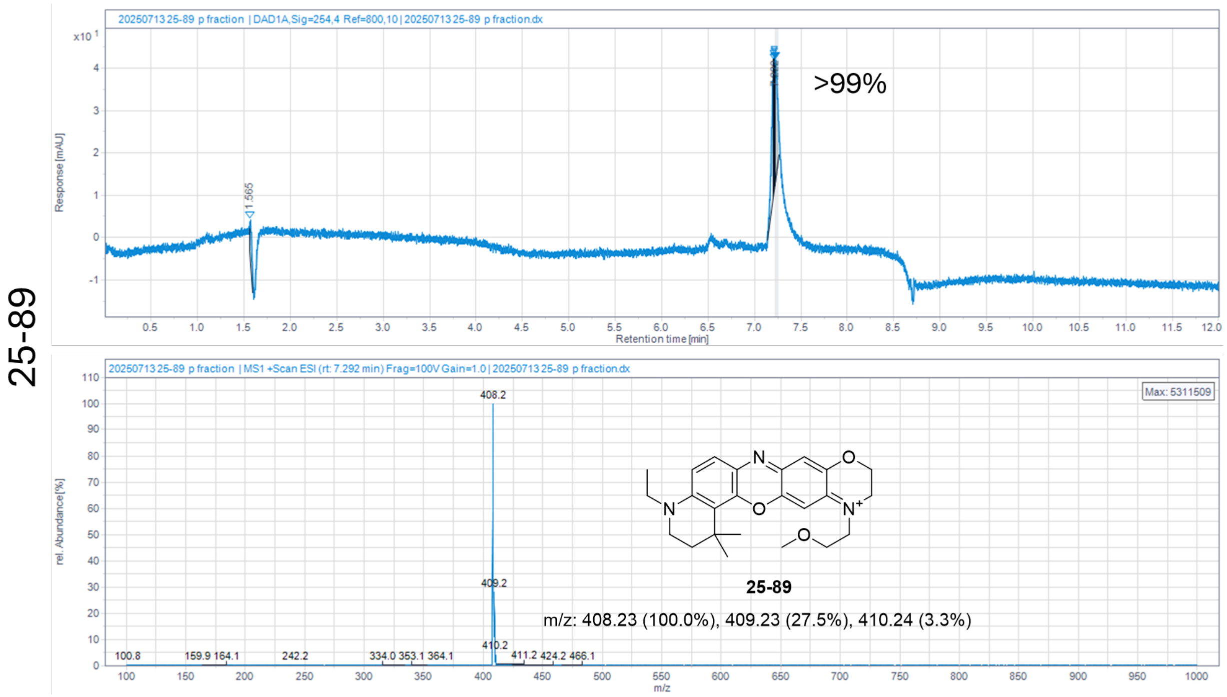


**Figure S31: HPLC-MS and purity analysis data for compound 25-89**

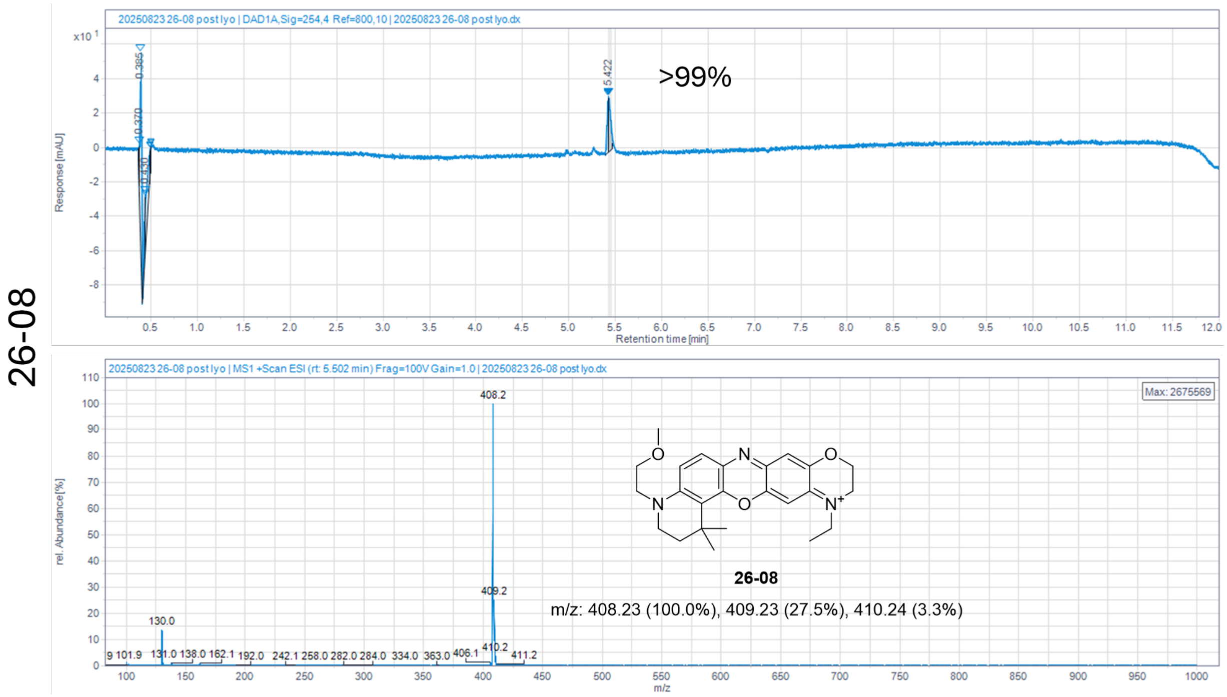


**Figure S32: HPLC-MS and purity analysis data for compound 26-08**

26-09

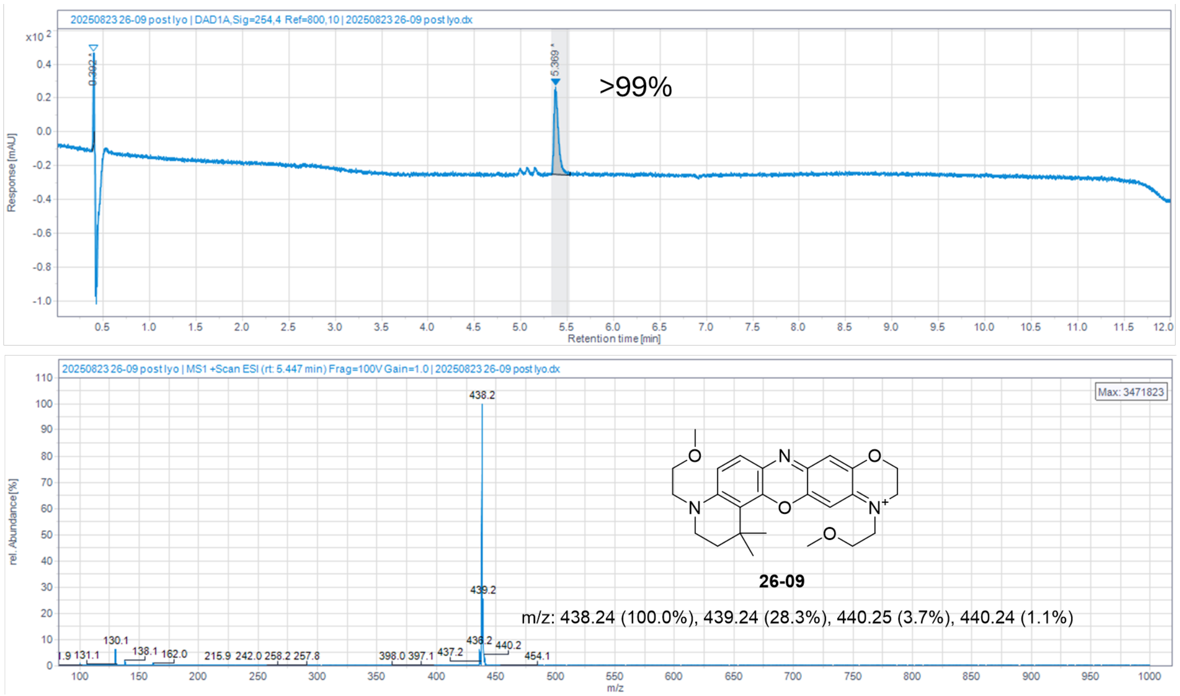


**Figure S33: HPLC-MS and purity analysis data for compound 26-09**

**Figure S16-33: HPLC-MS and purity analysis of oxazine dyes.** High performance liquid chromatography-mass spectrometry (HPLC-MS) was used to characterize the structure and purity of each fluorophore; absorbance at 254 nm and mass-to-charge ratio are presented for each fluorophore. Compounds with trace impurities were analyzed based on area under the curve analysis of the 254 nm absorbance, and a percent purity was determined.

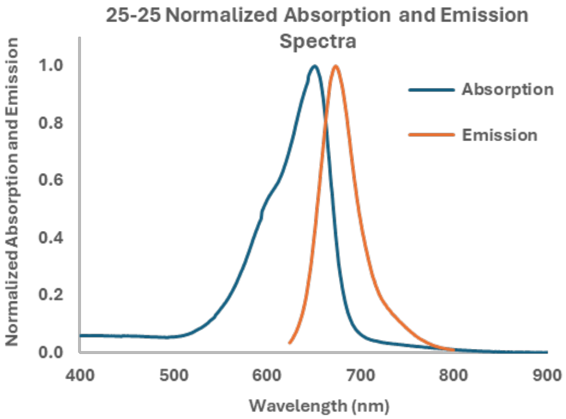


**Figure S34: Normalized Absorption and Emission Spectra for 25-25**

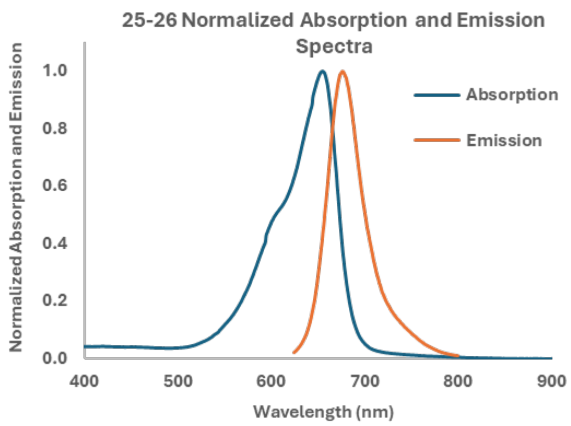


**Figure S35: Normalized Absorption and Emission Spectra for 25-26**

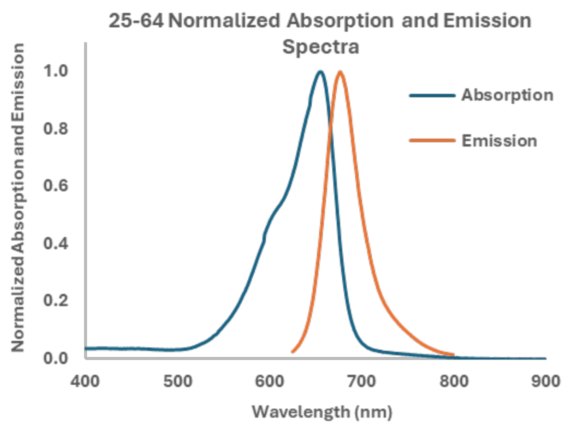


**Figure S36: Normalized Absorption and Emission Spectra for 25-64**

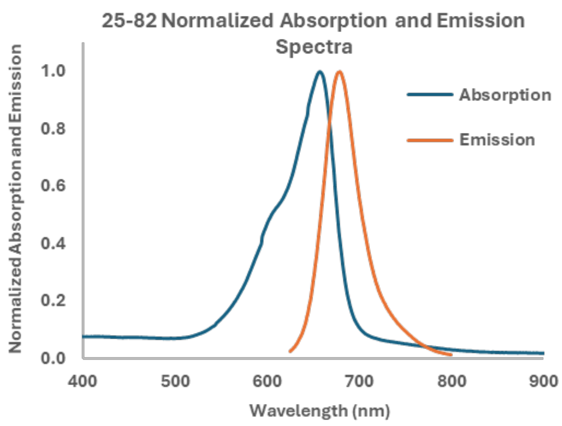


**Figure S37: Normalized Absorption and Emission Spectra for 25-82**

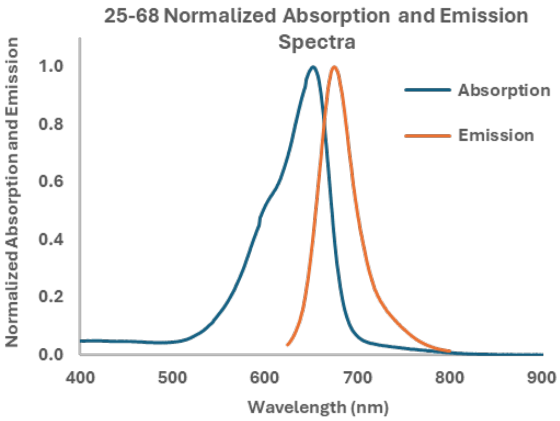


**Figure S38: Normalized Absorption and Emission Spectra for 25-68**

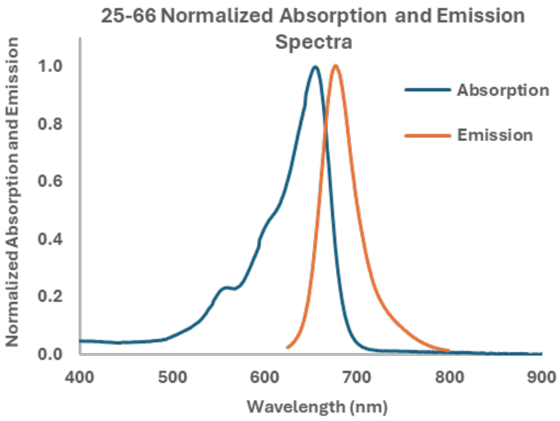


**Figure S39: Normalized Absorption and Emission Spectra for 25-66**

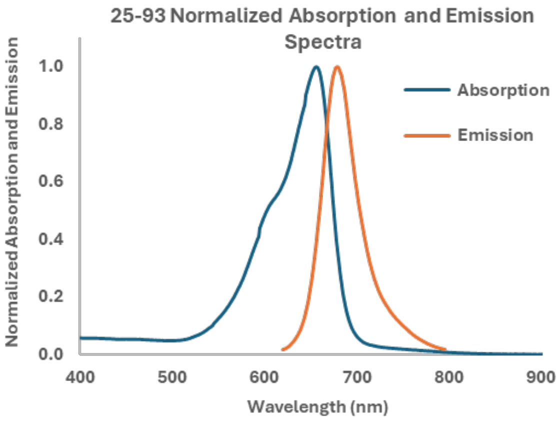


**Figure S40: Normalized Absorption and Emission Spectra for 25-93**

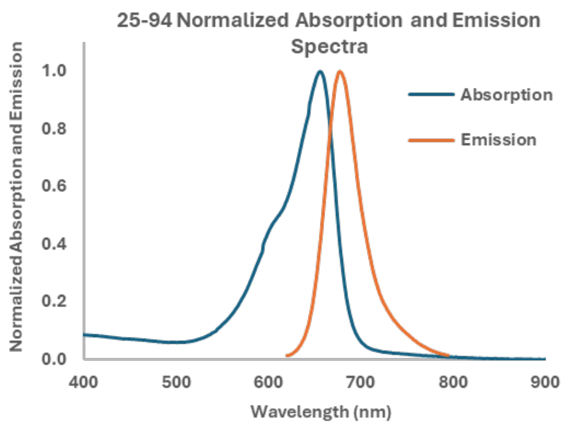


**Figure S41: Normalized Absorption and Emission Spectra for 25-94**

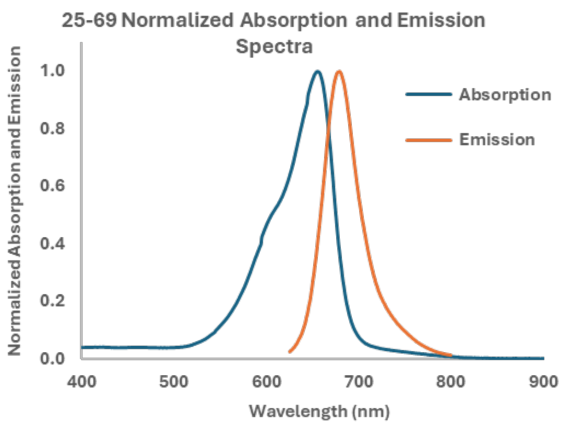


**Figure S42: Normalized Absorption and Emission Spectra for 25-69**

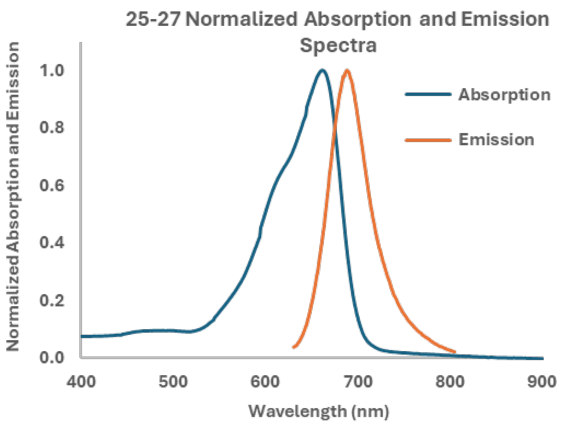


**Figure S43: Normalized Absorption and Emission Spectra for 25-27**

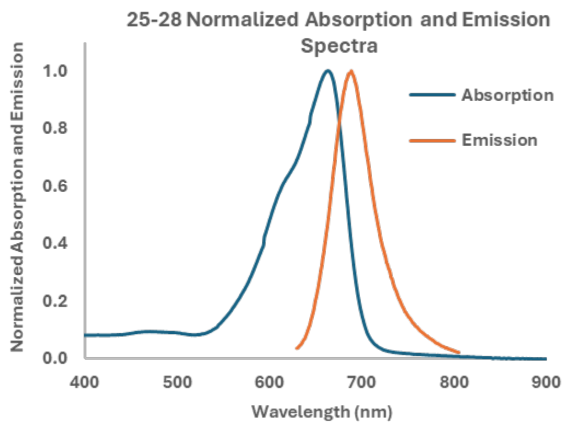


**Figure S44: Normalized Absorption and Emission Spectra for 25-28**

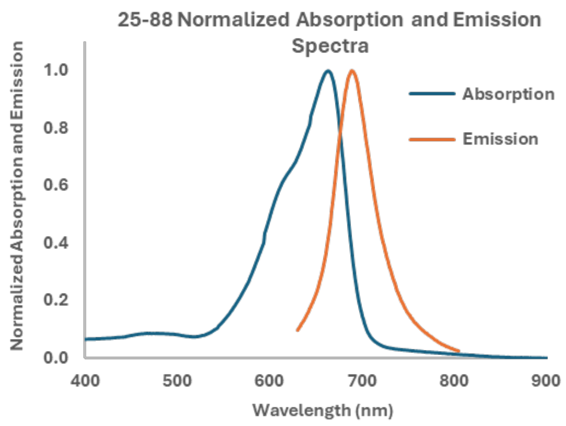


**Figure S45: Normalized Absorption and Emission Spectra for 25-88**

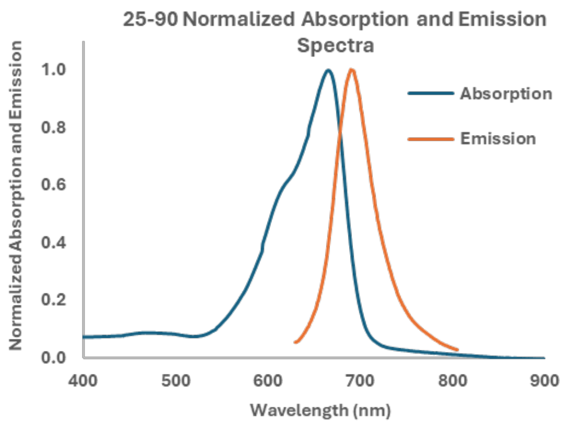


**Figure S46: Normalized Absorption and Emission Spectra for 25-90**

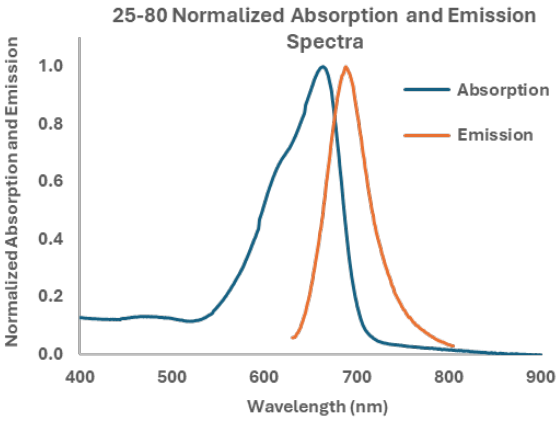


**Figure S47: Normalized Absorption and Emission Spectra for 25-80**

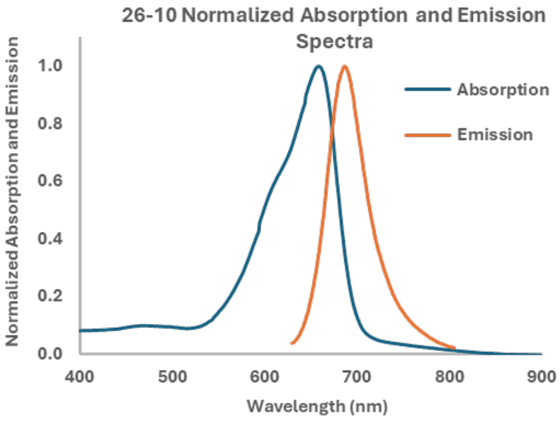


**Figure S48: Normalized Absorption and Emission Spectra for 26-10**

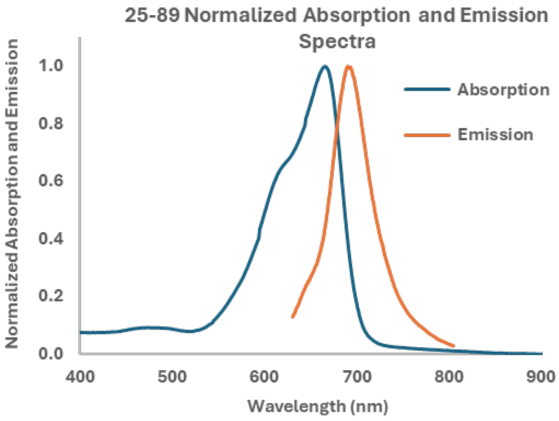


**Figure S49: Normalized Absorption and Emission Spectra for 25-89**

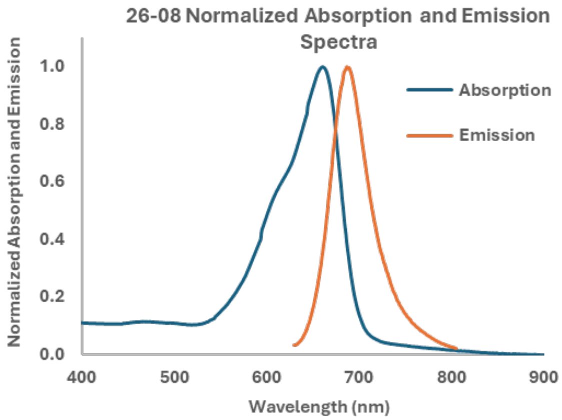


**Figure S50: Normalized Absorption and Emission Spectra for 26-08**

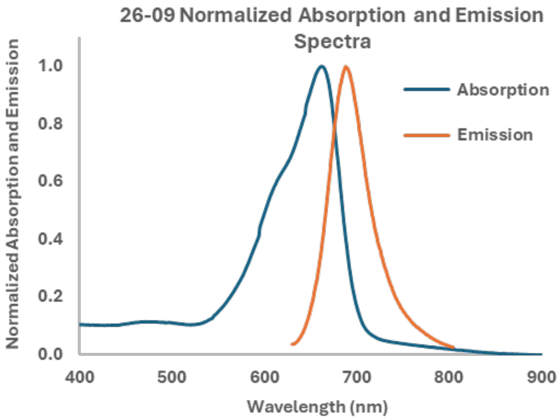


**Figure S51: Normalized Absorption and Emission Spectra for 26-09**

**Figure S34-51: NIR oxazine dye spectra.** Absorbance and emission spectra of oxazine fluorophore collected in Dulbecco's phosphate buffered saline (DPBS) normalized to highest absorption and emission value respectively. Displayed spectra are averages of multiple samples (n=3) of absorption and emission spectra for each fluorophore.
